# PhytClust: Efficient and Optimal Monophyletic Partitioning of Rooted Phylogenetic Trees

**DOI:** 10.64898/2025.12.11.693738

**Authors:** Katyayni Ganesan, Elisa Billard, Tom L. Kaufmann, Cody B Strange, Maja C. Cwikla, Adrian M. Altenhoff, Christophe Dessimoz, Roland F. Schwarz

## Abstract

Phylogenetic trees play a fundamental role in elucidating evolutionary relationships among taxa. Clustering taxa remains a major challenge across diverse biological domains such as cancer genomics, microbial systematics, and phylogenomics. Several methods partition taxa in phylogenetic trees into clusters, but these approaches face key limitations. Many rely on user-defined distance thresholds, whose choice often depends on prior knowledge and may be difficult to justify consistently across datasets. More broadly, existing methods often differ in how clusters are defined and commonly rely on heuristics or user-specified criteria to make the search space tractable for large trees.

Here, we present PhytClust, a threshold-free algorithm that partitions trees into monophyletic subtrees (clusters) by identifying groups of taxa with low within-cluster dispersion. For a fixed number of clusters, PhytClust finds the exact global optimum under this objective and then selects the optimal number of clusters using a cluster-validity index. The resulting partitions are reproducible and reflect both tree topology and branch lengths.

In simulated datasets, PhytClust outperforms existing methods in both speed and accuracy and scales to trees with more than a hundred thousand taxa. We apply PhytClust across cancer genomics, avian phylogenomics, bacterial and archaea phylogenetics, and plant genomics to demonstrate PhytClust’s varied applicability. By providing a standardized method for taxa clustering within phylogenetic trees, PhytClust yields reproducible, optimal and computationally efficient clusters.

## Introduction

Phylogenetic trees are the primary means of quantitatively analyzing relationships among taxa and underpin many areas of biological research^1^. Phylogenetic inference collapses large genetic datasets into a coherent evolutionary narrative^2^, enabling the identification of branching patterns and genealogical hierarchies, reconstruction of ancestral states, and tests of evolutionary hypotheses^3,4^. Phylogenies are widely used to track pathogen trajectories, dissect tumour evolution, reveal trait diversification across species and inform bioconservation strategies ^5–7^.

A recurring challenge in these applications is identifying robust subgroups of leaf nodes within a phylogeny. These subgroups or clusters are typically sets of closely related taxa that are meaningfully distinct from other taxa in the tree. In species evolution, such clusters can naturally emerge as the result of external factors such as changing environmental conditions leading to adaptive radiation or a change in the interplay of anagenesis and cladogenesis, or genetic events such as gene duplications **(Fig. 1a)**. In systematics, clusters are often aligned with taxonomic ranks, but manual curation is difficult for closely related species lacking clear synapomorphies and can yield uneven diversity across equivalent ranks^8^. Additionally, as new species are added to the tree of life, classification becomes more difficult. For example, in the recently discovered *Asgard* archaea, subfamilies were curated manually by the researchers, and these classifications have to be re-evaluated any time new species are added to the tree^9– 11^. Similarly, in cancer research, tumour phylogenies are reconstructed based on somatic mutations found in cancer cells and *subclones*, i.e. cancer cells with similar, not necessarily identical, genetic profiles, are often identified manually or using arbitrary thresholds^12–14^. Analogous issues arise when defining *operational taxonomic units* in microbiome research or delineating viral lineages in epidemiology^15^.

**Fig. 1:**
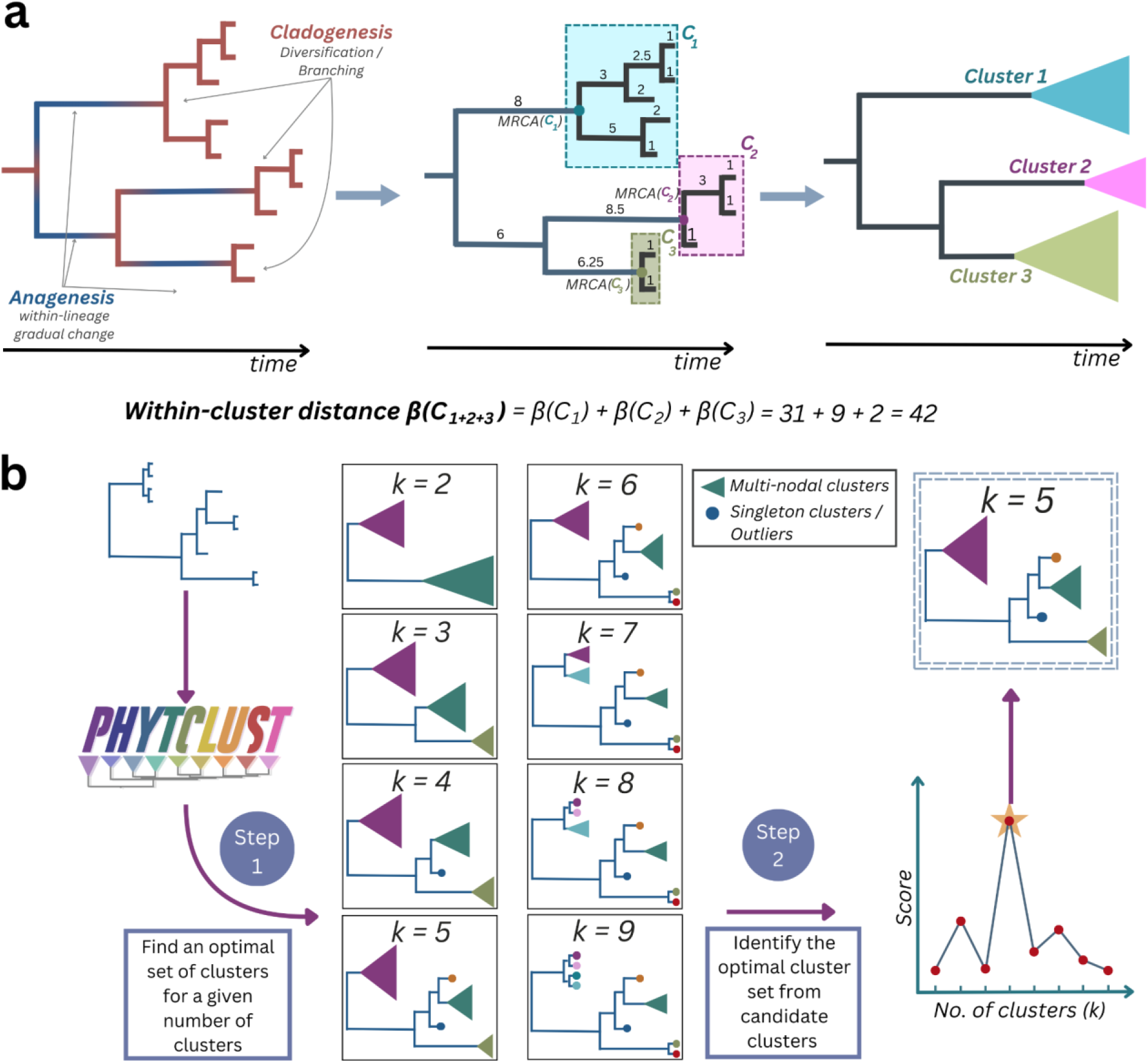
(a) An illustrative example of a phylogenetic tree in which both anagenetic and cladogenetic processes are partitioned into three family-level clusters (C1, C2, C3). Clusters are defined by minimizing the cumulative within-cluster distance β, calculated by summing the distances from each taxon to its most recent common ancestor (MRCA) within the cluster. (b) PhytClust workflow: Given a phylogenetic tree, PhytClust first computes the optimal clustering solution for every possible number of clusters k ∈ [1, N], where N is the number of leaves in the tree, including the trivial k = 1 and k = N cases. It then evaluates the non-trivial candidates k ∈ [2, N − 1] using the cluster-quality index and selects the optimal solution k*. The trivial cases are omitted from the schematic for simplicity.

Systematic, objective, and reproducible clustering directly on tree geometry therefore remains an essential goal. On rooted trees, clusters can be defined by a vertical cut at a fixed distance from the root or tip, but this is most natural for ultrametric trees, where all leaves lie at the same root-to-tip distance.^16^. In non-ultrametric trees, however, no simple solution exists, and many algorithms depend on manual intervention or user-defined thresholds that can limit reproducibility and complicate comparison across studies^16,17^.

Existing tools illustrate these trade-offs. Early approaches for transmission clusters, such as ClusterPicker and PhyloPart, define a monophyletic clade as a cluster if within-clade patristic distances fall below a user-specified bound (maximum for ClusterPicker; median for PhyloPart)^18,19^. Later methods, including TreeCluster, enforce bounds such as maximum diameter or total within-cluster branch length on each cluster^16^. PhyCLIP views each internal-node subtree as a candidate cluster and uses size and diversity constraints and statistical tests within an integer-linear program^17^. Across these strategies, user-specified parameters remain central, complicating reproducibility and cross-study comparison. At the same time, thresholds often serve as proxies that cap computational burden in the vast space of possible tree partitions.

A more recent approach, AutoPhy^20^, reduces threshold dependence by clustering in an embedded distance space. It projects patristic distances using Uniform Manifold Approximation and Projection^21^, fits a Gaussian mixture model^22^ with the number of components chosen by the Bayesian information criterion^23^, and then splits provisional clusters into monophyletic clades when necessary. While this reduces explicit cutoffs, runtime can vary with topology and tree size (for example, over 100 minutes on a ∼3,900-tip SARS-CoV-2 tree)^20^. Collectively, these limitations motivate methods that (i) operate directly on phylogenies, (ii) avoid user-defined thresholds, (iii) are exact for a given model, and (iv) scale to large trees.

To address these limitations, we have developed PhytClust. PhytClust divides any rooted phylogenetic tree into disjoint sets of monophyletic clusters (clades) based on a well-defined mathematical criterion of minimizing within-cluster dispersion, defined as the sum of tip-to-MRCA distances within clusters. PhytClust implements an exact dynamic programming (DP) algorithm that computes the global optimum for any specified number of clusters *k*. PhytClust then uses a tree-specific goodness-of-fit quality index to select an optimal non-trivial clustering resolution from candidate values *k* ∈ [2, *N* − 1], while computing optimal partitions for all *k* ∈ [1, *N*]. On simulated data, PhytClust outperforms existing methods while remaining competitive in runtime. We show PhytClust’s applicability on real-world trees including single-cell cancer phylogenies, species trees from various biological domains including bacteria, archaea, plants and birds, and on gene trees where it accurately identifies gene duplication events. By anchoring clusters to monophyletic clades and optimizing a transparent, branch-length–based objective, PhytClust yields standardized, reproducible partitions that respect ancestry and minimize within-cluster divergence.

## Results

### PhytClust delineates monophyletic clades in rooted phylogenetic trees

We formalize clustering on rooted phylogenies as the problem of partitioning the leaves into *k* disjoint monophyletic clusters. Requiring clusters to be monophyletic ensures that each cluster corresponds to a clade defined by common ancestry, rather than being defined solely by a user-specified distance cutoff. Similar to clustering concepts in Euclidean spaces, we strive for clusters of taxa that are closely related within the clusters, while remaining relatively distant between the clusters. Thus, for any fixed number of clusters *k*, we define the optimal partition as the one that minimizes the within-cluster dispersion *β* **(Fig. 1a)**. Here, compactness is measured as the sum of tip-to-MRCA distances within each cluster, treating the MRCA as a cluster-specific centroid. Minimizing within-cluster dispersion simultaneously increases between-cluster separation, thereby indirectly maximizing separation between clusters **(Supplementary Proof. A)**.

For a given value of *k*, identifying the set of monophyletic clusters that minimize *β* through exhaustive enumeration is only feasible for small trees **(see Methods; Supplementary Fig. 1a)**. Because empirical phylogenies typically fall between the two topological extremes of fully imbalanced caterpillar trees and perfectly balanced binary trees^24^, their numbers of possible monophyletic partitions are also expected to fall between the corresponding bounds **(Eq. (6))**. To efficiently obtain optimal partitions for each possible number of clusters *k*, we make use of the tree’s natural hierarchical structure. We recursively divide the tree into successively smaller subtrees and construct the global solution from the solutions to these subproblems using dynamic programming **(Fig. 2a)**. Briefly, for each node, we compute the minimum *β* attainable with every possible number of clusters within that node’s subtree. These intermediate results are stored so that each subproblem is solved only once. By combining the stored results from descendant subtrees, we iteratively determine the optimal partitions for each parent node, continuing upward through the tree until the global optimum at the root is obtained. Multiplying all branch lengths by a positive constant leaves both the optimal partition for any fixed *k* and the selected *k* unchanged, since all costs scale equally, making PhytClust invariant to global rescaling.

**Fig 2:**
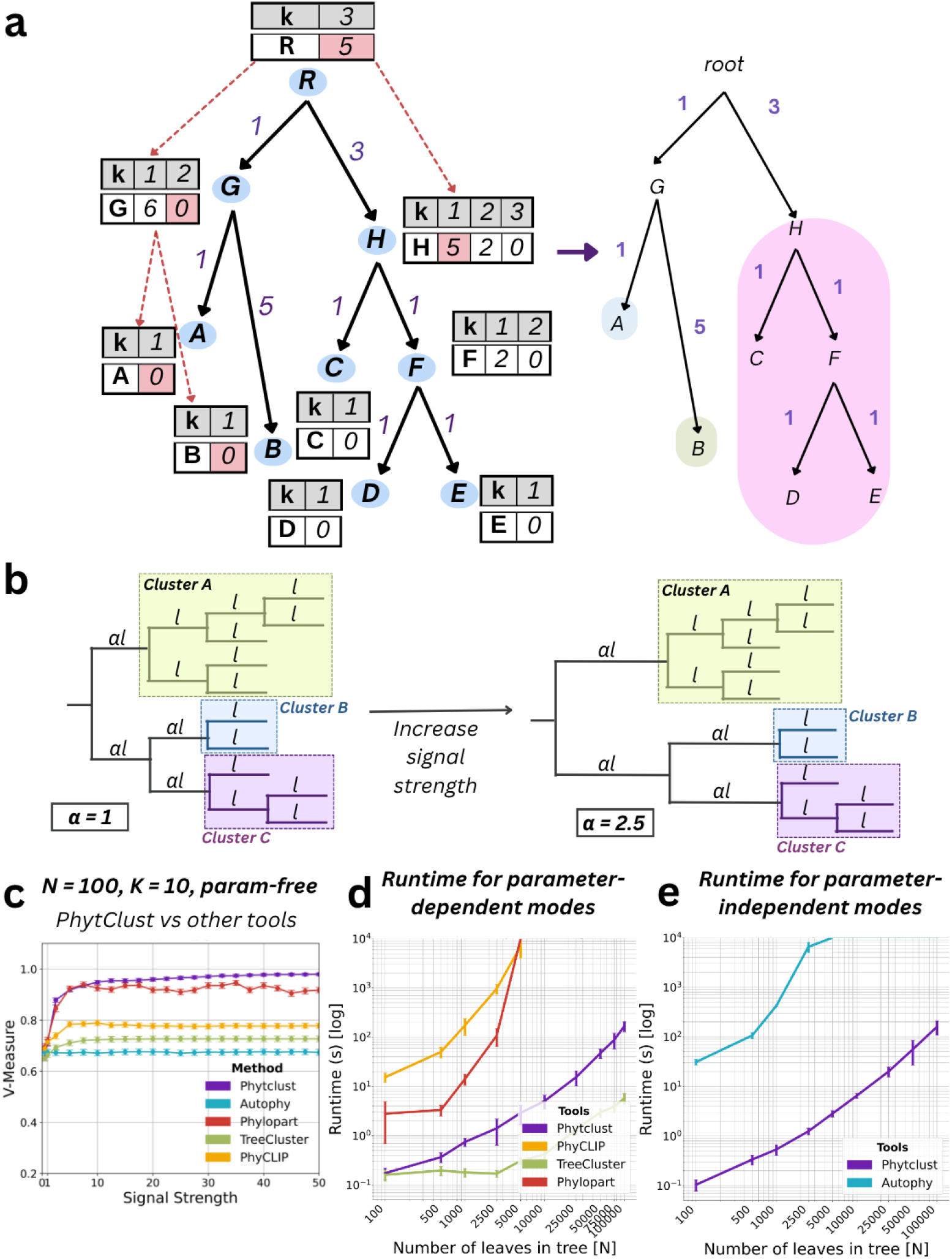
(a) Example dynamic programming table for a sample tree. For k = 3, the entry at the root points to child H as one cluster and to child G as split into subclusters. The resulting optimal set of clusters that minimizes β is [A], [B], [C, D, E]. (b) Simulation setup: Random trees were generated with initial branch length of l = 1 and α = 1, and k random monophyletic target clades were assigned. Signal strength α, defined as the ratio of average between-to within-cluster dispersion, was increased on extra-cluster branches. (c) Simulation results reporting V-Measure for PhytClust, AutoPhy, PhyCLIP, TreeCluster and PhyloPart of trees with N =100, k = 10. (d) Runtime comparison of software with a parameter-dependent mode (PhytClust, TreeCluster, PhyloPart, PhyCLIP) (e) Runtime comparison of software with a parameter-independent mode (PhytClust, AutoPhy).

From the globally optimal partitions for each possible number of clusters *k*, PhytClust uses a cluster-quality index to select the optimal number of clusters (*k*)* combining two known indices—the Calinski–Harabasz (CH) and the elbow index (EL1)—and adapting them for rooted trees **(see Methods, Fig 1b)**^25,26^. The cluster-quality index enables the selection of partitions that balance compactness within clusters and separation between clusters. The total branch-length cost *β*(*k*) computed by our dynamic programming procedure thereby corresponds to the within-cluster term *W*(*k*) of the CH index **(see Methods and Supplementary Proof A)**. This relationship allows CH to be computed directly from *β*(*k*) without additional traversals over the tree, limiting the computational complexity and allowing for efficient selection of the optimal clustering resolution *k*\*. The EL1 index further highlights the point of diminishing returns in *β*(*k*), where additional clusters provide little improvement in within-cluster compactness **(Methods, Supplementary Fig. 1d)**. PhytClust computes optimal partitions for all *k ϵ* [1, *N*], including the trivial cases *k* = 1 and *k* = *N*, but score-based selection of the optimal resolution is restricted to the non-trivial range *k ϵ* [2, *N* − 1].

Despite this, selecting a single optimal global resolution *k\** remains a heuristic and depends on the goals of the analysis. Real-world phylogenies often contain meaningful partitions at several hierarchical levels. In biological taxonomy, for example, organisms are organized into families, orders, and classes, each corresponding to a distinct level of evolutionary divergence. Similarly, in viral or microbial phylogenies, one may wish to identify both broad lineages and fine-grained strain clusters. Because a single *k* cannot capture all relevant levels at once, PhytClust extends beyond a single optimal resolution to report partitions across the tree’s hierarchy. To do this, we evaluate our quality index over *k ϵ* [2, *N* − 1] using log-spaced bins **(Methods)** and identify local maxima that represent well-supported partitions at each clade level. The resulting family of partitions summarizes structure across the full range of levels in the tree and allows users to interpret clusters according to their specific analytical goals without manually specifying *k*. To further lower the barrier to exploration, PhytClust includes an interactive web interface through which users can visualise clustering solutions across resolutions, inspect individual partitions, and export figures; several visualisations in this manuscript were generated directly through this interface.

### PhytClust accurately recovers simulated target partitions

We evaluated PhytClust on simulated rooted binary trees with tunable signal strength *α ϵ* [1, 50], defined as the ratio of average between-to within-cluster dispersion. For each condition defined by a fixed number of taxa *N* and true clusters *k*, we generated 100 rooted binary topologies drawn uniformly from all possible trees^24^. We then selected *k* disjoint monophyletic target clades, assigned all branches an initial length of one, and increased the between-cluster separation by multiplying branches ancestral to each true clade by *α* **(Fig. 2b; Methods)**.

As a null baseline, we generated 100 random monophyletic *k*-partitions with *α* = 1 for each tree. We quantified agreement between inferred and true clusters using the V-measure (*V*)^27^, an external clustering metric ranging from 0 to 1, with higher values indicating better recovery of the reference partition. For these random partitions **(Supplementary Fig. 2a)**, the V-measure remained close to 0.5 across all values of *α*. This baseline is above zero because the random partitions are constrained to be monophyletic on the same underlying tree and therefore can show non-trivial agreement with the reference partition even without branch-length signal. In contrast, the V-Measure of PhytClust increased steadily with signal strength. For *k* = 10 the median V-measure exceeded 0.90 at *α* = 15 and reached *V* = 1.00 for all replicates once *α* > 45 **(Supplementary Fig. 2e)**. When *α* = 1, within- and between-cluster distances are indistinguishable, so performance approaches a monophyletic-random baseline determined by the tree topology and cluster size distribution **(Supplementary Fig. 1c, 1d)**.

When allowing PhytClust to automatically select the number of clusters (*k*)* using our cluster-quality index, we compared its performance with PhyCLIP, TreeCluster, PhyloPart, and AutoPhy **(Methods)**. TreeCluster and PhyloPart each require a single user-defined distance threshold that determines how partitions are made, and their performance depends strongly on this choice. To ensure a fair comparison, we performed a grid search over a wide range of thresholds and reported results that maximized the silhouette score^28^ **(Fig. 2c; Methods)**, which favors high between-cluster separation relative to within-cluster cohesion. PhyCLIP was run across a broad range of its parameters, and its intermediate-resolution mode was used to automatically select one clustering from the multiple solutions it generates.

At low signal strength (*α* = 1), all methods achieved similar V-measure values around 0.65, reflecting the lack of separation between true clades **(Fig. 2c)**. As signal increased, AutoPhy remained near *V* = 0.66, TreeCluster and PhyCLIP plateaued around *V* = 0.70 and *V* = 0.80 by *α* = 10, and PhyloPart reached about *V* = 0.90 at *α* = 7.5 with higher variability. PhytClust exceeded *V* = 0.95 already at *α* = 15 and approached *V* = 0.97 for stronger signals, achieving the highest V-measure among the methods compared, even at moderate signal strengths. The same trend was observed for *k* = 5 and *k* = 20 **(Supplementary Fig. 2c)**. For *N* = 100 and *k* = 10, qualitative examples show clear under-clustering at low signal, partial recovery at moderate signal, and exact recovery at high signal **(Supplementary Fig. 1g, 3-5)**. Moreover, we examined whether PhytClust’s performance depended on backbone-tree balance and cluster-size inequality. Of these two factors, only cluster-size inequality, quantified by the Gini index^29^, where higher values indicate greater inequality in cluster sizes, showed a clear effect **(Supplementary Fig. 2d)**.

We next benchmarked runtime across random bifurcating trees containing up to *N* = 10^5^ tips (100 replicates per tree size). When using fixed thresholds for PhyloPart, TreeCluster, and PhyCLIP, or a fixed *k* for PhytClust, TreeCluster was fastest and PhytClust a close second **(Fig. 2d)**. On trees with 10^5^ tips, TreeCluster completed in under 10 seconds, while PhytClust required approximately 100 seconds, remaining practical for trees of 10^5^ tips. In threshold-free comparisons, we included only PhytClust and AutoPhy, since these methods do not rely on any user-defined cut-offs or parameter grids. Other tools were excluded from this comparison because their runtimes depend strongly on the range of parameters supplied. In this setting, PhytClust ran faster than AutoPhy **(Fig. 2e)**. While the worst-case theoretical time complexity of PhytClust is O(*N*^*2*^), power-law fit to measured runtimes yielded an average exponent of ∼1.38 across random trees, indicating subquadratic scaling in practice, though the exponent varies with tree topology between the linear (caterpillar) and quadratic (balanced) extremes **(Supplementary Fig. 14d)**. On Genome Taxonomy Database^30^ phylogenies, PhytClust clustered the Archaea (5,869 taxa) in 5 s and the Bacteria (107,235 taxa) in 9min 11s, demonstrating scalability to genome-scale datasets.

Taken together, these results show that PhytClust combines high accuracy with practical runtime performance, remaining efficient even for trees containing more than 100,000 taxa.

### Empirical phylogenies span the simulated signal range

To assess whether the signal-strength range identified in our simulations is also relevant in empirical data, we next estimated *α* as a descriptive summary statistic for a set of real phylogenies. Here, *α* denotes the ratio of average extra-cluster to average within-cluster dispersion with respect to curated reference clades (Methods).

Using phylogenies from Nextstrain^31,32^, we computed *α* for established clades in dengue, monkeypox, tick-borne encephalitis^33^ and tuberculosis, which were 85.5, 25.6, 13.1, and 6.8 respectively. For an independent coronavirus phylogeny^34^ delineating seven major groups, we obtained *α* = 62.5. Together, these values span a broad portion of the simulated range (*α ϵ* [1, 50]) and show that empirical trees can include both high-signal and substantially lower-signal cases **(Methods)**.

The tuberculosis phylogeny is particularly informative because it lies near the low end of the tested regime. As a slowly evolving clonal bacterium^35–37^ sampled densely within major lineages, *M. tuberculosis* shows only modest branch-length separation among major lineages relative to within-lineage diversity. Even in this weak-signal setting, the PhytClust partition at fixed *k* = 11closely matched the 11 major annotated lineages (*V* = 0.98), and *k* = 11 also appeared among the five strongest candidate resolutions in the validity-index scores **(Supplementary Fig. 9)**.

Performance was similarly strong across the viral datasets. In the coronavirus phylogeny, PhytClust recovered all seven groups as the automatically selected solution *k\** **(Supplementary Fig. 6a)**. In dengue, PhytClust selected *k*\* = 5, where the additional fifth cluster corresponded to the deeply diverged sylvatic DENV2 clade^38,39^, yielding near-complete agreement with serotype-level annotation (*V* = 0.99; **Supplementary Fig. 6b**). At finer resolution, the fixed-*k* solution with *k* = 20 also agreed closely with the 20 annotated dengue genotypes (*V* = 0.95; **Supplementary Fig. 6b**). For tick-borne encephalitis virus, the automatically selected *k\** matched the standard clades^33^ except for one split, whereas fixing *k* = 8 yielded exact concordance **(Supplementary Fig. 8)**. For monkeypox, *k\** = 3 merged Clades IIa and IIb into a single cluster, consistent with their relatively modest branch-length separation, while setting *k* = 4 recovered the canonical four **(Supplementary Fig. 7)**. For comparison, AutoPhy was also run on these empirical phylogenies, but feasible runtimes were obtained only for the monkeypox and tick-borne encephalitis datasets **(Supplementary Figs. 7-8)**.

Overall, these analyses show that the signal-strength regime explored in simulation is realistic for both viral and non-viral phylogenies. Real datasets can range from highly separated clade structure to much weaker branch-length contrast, yet PhytClust remains able to recover biologically meaningful partitions across this spectrum.

### PhytClust identifies meaningful subclones in cancer trees

Cancer cells evolve through point mutations and karyotypic changes such as copy-number (CN) alterations and whole-genome doubling (WGD)^40^, generating intra-tumour heterogeneity that complicates treatment. A common challenge in cancer genomics studies is defining subclones in cancer trees reconstructed from single-cell or multi-region bulk sequencing datasets, a task that is still often performed manually^13,14^. Here we instead define subclones objectively using PhytClust.

As an illustrative example, we re-analysed a single-cell phylogeny of a case of high-grade serous ovarian cancer from the SPECTRUM cohort^41^. In patient OV-025, PhytClust identified 14 clusters in total (*α* = 1.8). For clarity, we focus here on eight major subclones **(Fig. 3)**, with six additional clusters represented by fewer than 12 cells each (≈0.01% of total). PhytClust assigned the two key lineages reported by McPherson et al., which had independently acquired two WGD events each, to distinct subclones 6 and 8 **(Fig. 3, Supplementary Fig. 10)**. The other six major subclones each harbor a single WGD event, as previously shown, and exhibit distinct allele-specific CN alterations that clearly separate them from the other subclones. For example, subclone 7 lacks the high-level gains of its sister subclone 8 (two WGDs), instead showing lower copies across chromosome arms 12q and 13q and unique loss of heterozygosity (LOH) on 3q and 6q. Subclone 6 carries arm-level amplifications on 3q and 7q with a total copy number of 9 together with an LOH event on 14q. Even closely related subclones differ in clonality of shared events, for example, on chr1:77–88.5 Mb the allele-specific (3,2) gain occurs in ∼90% of subclone 1 cells vs ∼56% in subclone 2. Similarly, on chr7:147–149.5 Mb a combined amplification / LOH event with alleles-specific copy number (7,0) occurs in ∼88% vs ∼57% of cells.

**Fig 3:**
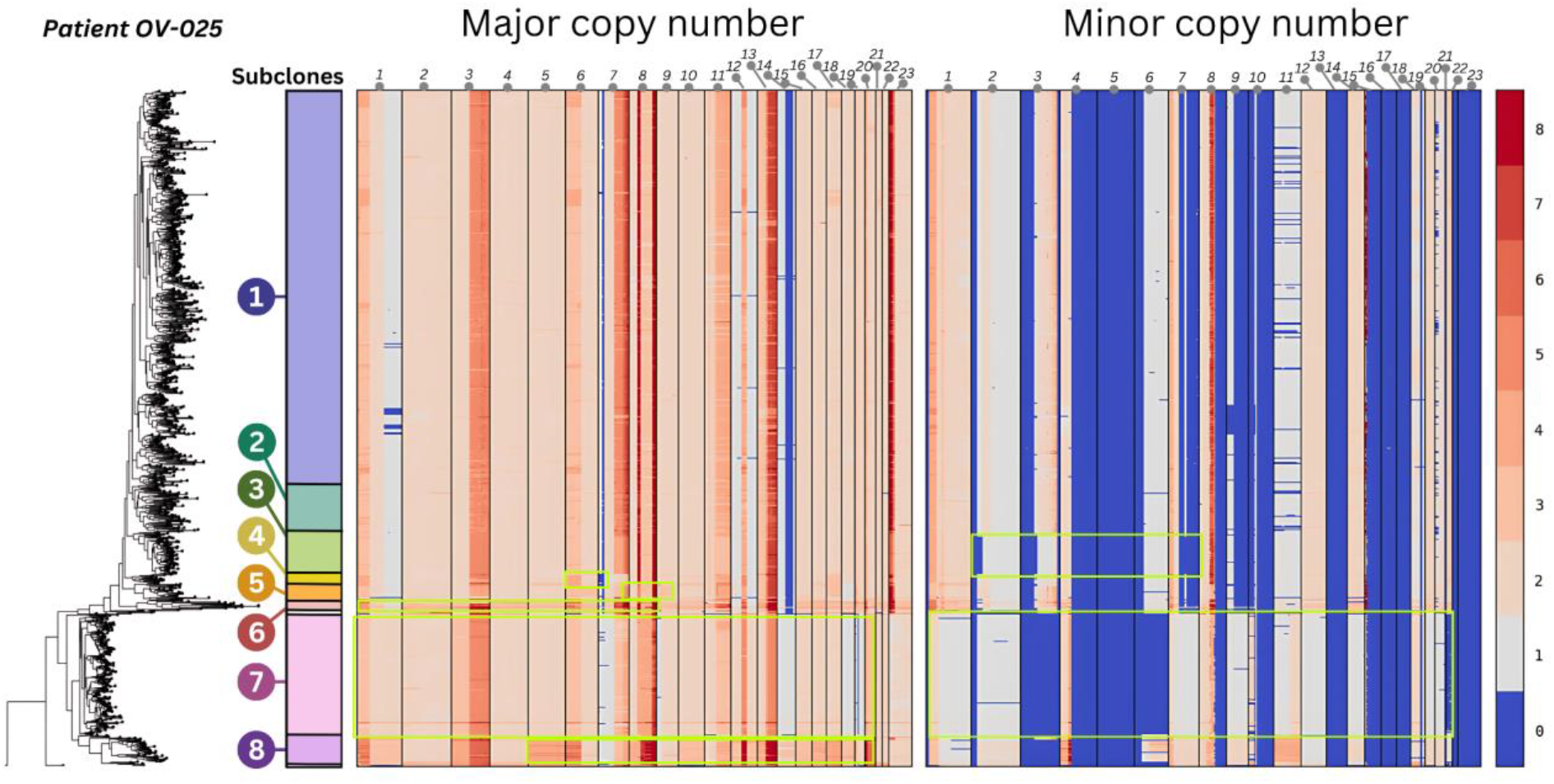
Phylogenetic tree of patient OV-025 with PhytClust-defined subclones and their associated copy-number profiles. The tree shows the OV-025 phylogeny, with the 8 major PhytClust subclones highlighted. Adjacent genome-wide allele-specific copy-number (CN) profiles summarize recurrent CN states within each highlighted subclone and illustrate the CN differences used for biological interpretation in the text. PhytClust identified 14 clusters in total; the remaining 6 are very small terminal clusters (<12 cells each) that are retained in the phylogeny but are not discussed as separate major subclones in the main CN summary. The color bar at right shows the copy-number scale.

We also reanalysed a breast cancer phylogeny (TN-2) originally studied by Minussi et al.^13^, with the tree reconstructed using MEDICC2^42^. Using the number of superclones (k = 6) and subclones (k = 10) reported by Minussi et al. and Kaufmann et al. as fixed inputs, PhytClust identified strictly monophyletic partitions at both resolutions **(Supplementary Fig. 11)**. Several of the original manually defined groups were not strictly monophyletic, mixing lineages with divergent evolutionary histories. By contrast, PhytClust’s clusters were evolutionarily coherent by construction and each supported by distinct allele-specific CN features, demonstrating that greater evolutionary homogeneity can be achieved without sacrificing biological interpretability.

Together, these results show that PhytClust yields reproducible, monophyletic subclone calls that respect evolutionary structure and surface biologically interpretable CN events, including independently doubled lineages, offering an objective alternative to manual or threshold-based subclonal delineation in cancer phylogenetics.

### PhytClust outlines hierarchical taxonomic groups in the Asgard phylogeny

The recent discovery of numerous Asgard archaeal lineages has reshaped our understanding of early eukaryotic evolution, but the pace of discovery has also made consistent taxonomic classification increasingly difficult^10,11,43^. As new metagenome-assembled genomes continue to expand this group, reproducible delineation of higher-order clades becomes challenging using traditional, threshold-based approaches^9,44,45^. To assess whether PhytClust can provide a stable, parameter-free alternative, we re-analysed the Asgard phylogeny from Liu et al. (2021b)^9^. Liu et al. assembled 209 single-copy *clusters of orthologous genes* (COGs), whose concatenated sequences were used to infer a maximum-likelihood tree. Liu et al. also delineated phylum-level clades by inspecting bootstrap support and gene-content similarities. Using that same tree as input, PhytClust selected *k\** = 12 as the optimal clustering resolution (*α* = 2.5) **(Fig. 4a)**.

**Fig 4:**
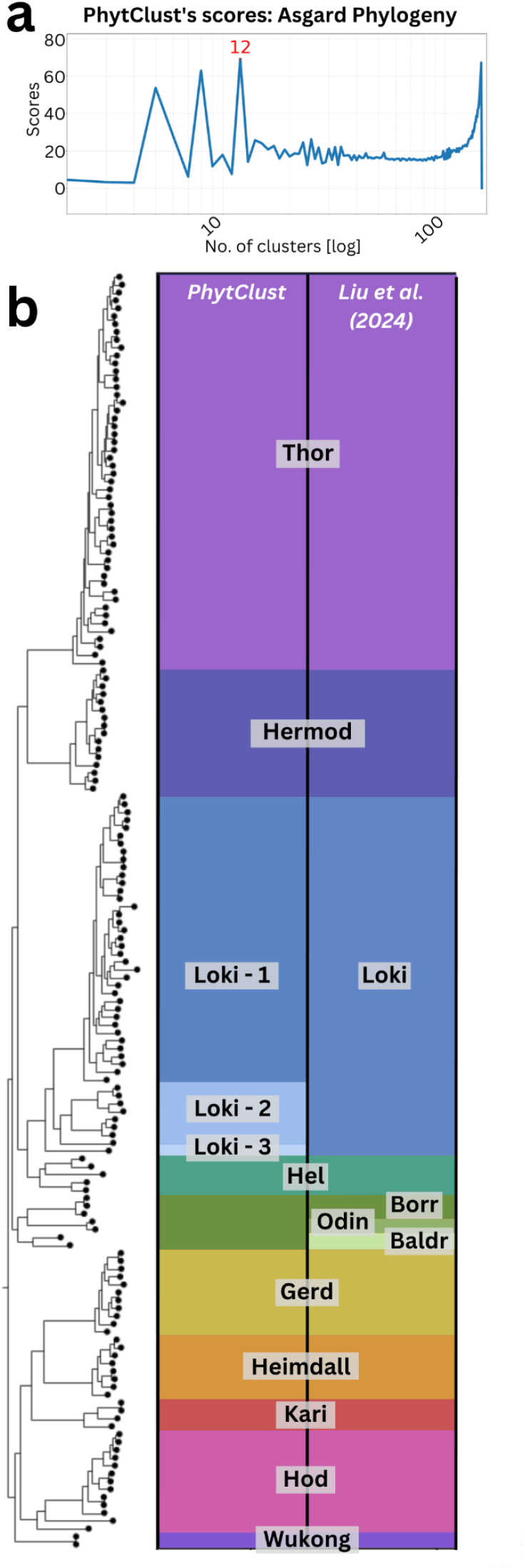
(a) PhytClust scores and selected solution at k* = 12 for the phylogenetic tree of Asgard archaea (b) Phylogenetic tree of the 12 sub-families of the Asgard family introduced by Liu et al. and the subfamilies proposed by PhytClust at k = 12.

PhytClust’s partitions showed extensive overlap with the classification of Liu et al. (*V* = 0.95), indicating that, on this phylogeny, tree topology and branch-length information alone largely recapitulate the expert-curated higher-level structure (**Fig. 4b)**. PhytClust subdivided Lokiarchaeota into three phylum-level clusters, while merging together Borrarchaeota, Baldrarchaeota and Odinarchaeota—consistent with the substantially greater taxonomic diversity of Lokiarchaeota relative to the latter groups combined. Notably, this finer subdivision receives independent support from the Genome Taxonomy Database: all species clusters in Loki-2 belong to a single family (SOKP01, order Signyarchaeales)^30^, consistent with PhytClust recognizing a coherent taxonomic unit within the broader Lokiarchaeota. Applied directly to the published phylogeny, PhytClust reconstructed nearly all of the higher-level groupings reported by Liu et al., without requiring additional clustering thresholds or external taxonomic annotations.

### PhytClust identifies common taxonomic groups in Avian phylogeny

Biological classification is inherently hierarchical, with monophyletic groups observable at multiple taxonomic levels—from species and genera to families and orders—each reflecting distinct evolutionary levels. Accurately capturing these nested relationships in large phylogenies remains a key challenge for reproducible taxonomy.

To assess how PhytClust recovers known taxonomic structure across multiple levels of divergence, we applied it to a phylogeny of 363 bird species encompassing 92% of recognized families **(Fig. 5)**^46^. The phylogeny, reconstructed by Stiller et al., was annotated according to established family- and order-level classifications^47,48^. To enable systematic comparison with this reference taxonomy, we stratified the tree into five granularity levels (CL1-CL5) from coarse to fine, and selected the best-supported local optimum within each level **(see Methods, Supplementary Fig. 12a)**.

**Fig 5:**
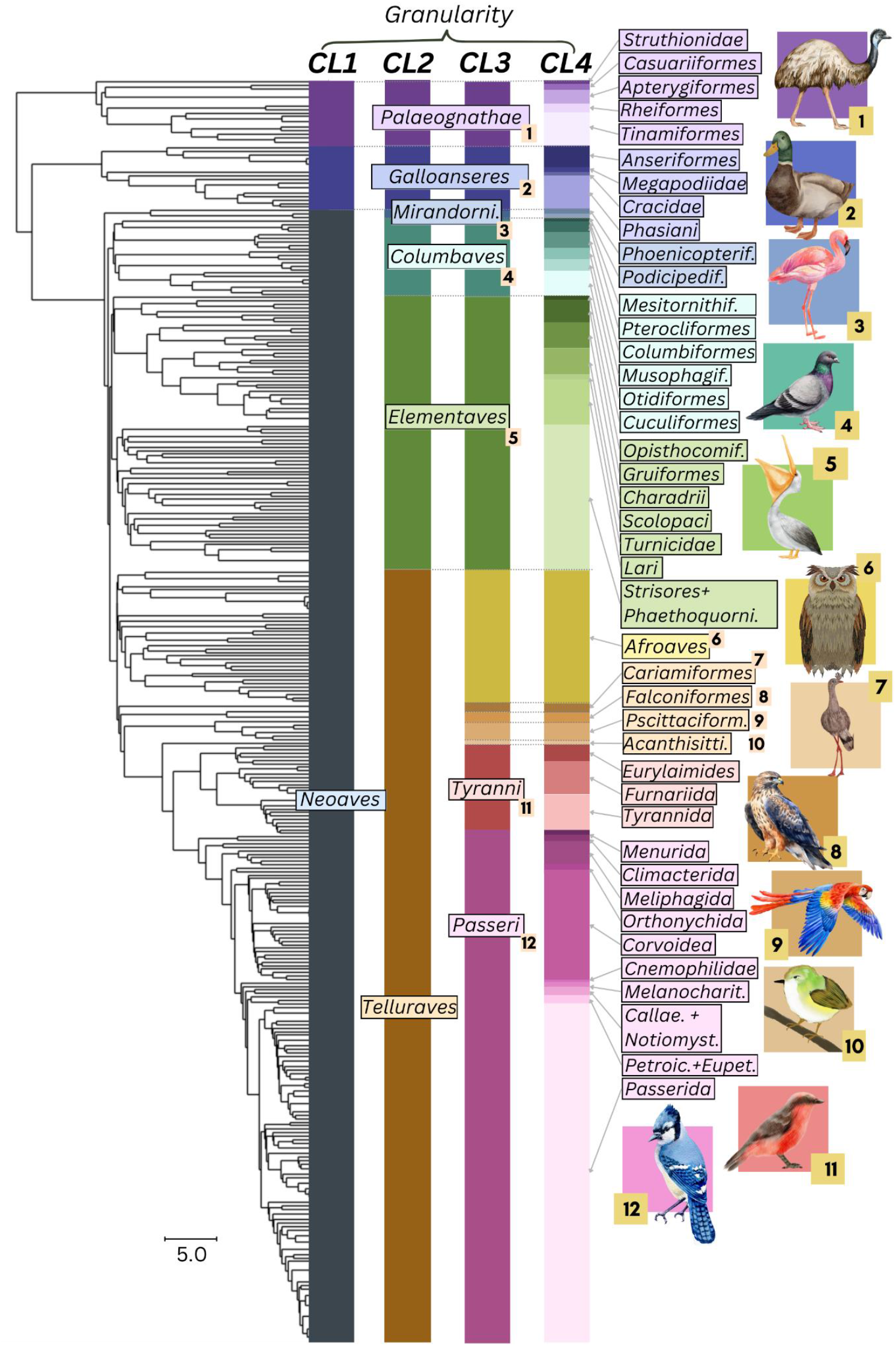
PhytClust was applied to the most comprehensive avian phylogeny available (from Stiller et al.^36^) and its scores were stratified into five clade levels (CLs; four shown). Clusters were labelled according to exact matches with existing taxonomic ranks or, when absent, the closest combination of ranks. Clusters from CL3 are numbered 1–12.

At the highest level (CL1), PhytClust accurately recovered the three major avian clades— Palaeognathae, Galloanserae, and Neoaves—demonstrating that the method captures broad evolutionary divisions consistent with classical taxonomy. The next level (CL2) produced five higher-order groups that aligned closely with published classifications, including the proposed Elementaves clade. We used the third level (CL3) for comparison with the published higher-level taxonomy because it provided the closest resolution to the corresponding reference grouping. CL3 (*α* = 0.9) refined these groups into twelve clusters and showed strong overall agreement with the reference classification (*V* = 0.85) **(Supplementary Fig. 12b)**. The main difference was that PhytClust split Telluraves rather than separating Columbaves and Elementaves, indicating that an objective branch-length-based partition can suggest a different but still biologically interpretable subdivision of this part of the tree. Moreover, PhytClust achieved a 15% reduction in cumulative within-cluster dispersion (reference: 2.3 × 10^4^ ; PhytClust: 1.9 × 10^4^), indicating greater internal compactness given our cluster-validity index objective.

At the fourth level (CL4), PhytClust identified 41 clusters with 10 singletons, compared with 53 clusters and 19 singletons in the reference taxonomy. This partition likewise had lower within-cluster dispersion under the same objective (reference: 1.5 × 10^4^; PhytClust: 1.3 × 10^4^). Most clusters aligned closely with existing classifications, and the observed differences were readily interpretable in phylogenetic terms. For example, PhytClust subdivided the diverse Passeriformes, consistent with their substantial internal diversity, while keeping the smaller Afroaves group intact.

Together, these results show that, on this avian phylogeny, PhytClust recovers hierarchical structure that overlaps extensively with established taxonomy across multiple levels of resolution. The inferred partitions are also more compact under the PhytClust branch-length-based objective than the corresponding rank-based reference groupings. These results support the utility of PhytClust as a reproducible tree-based framework for exploring taxonomic structure across resolutions, without requiring manually chosen clustering thresholds.

### Detecting duplication-driven clusters using PhytClust

Gene duplication is a major driver of evolutionary innovation^49^. Following duplication, one gene copy typically retains its ancestral function, while the other experiences relaxed selective constraints and may accumulate mutations^49,50^. This relaxed evolutionary pressure promotes functional divergence and is often reflected in accelerated sequence evolution, producing longer branches for duplicated genes compared to single-copy orthologs^49,50^.

These patterns provide a recognizable phylogenetic signal of post-duplication divergence. We hypothesized that such signals could be systematically detected using PhytClust, which clusters trees based solely on branch-length information and topology. Specifically, we tested whether PhytClust preferentially identifies clusters whose MRCA parent node is annotated as a duplication event.

We analysed 15,693 phylogenetic gene trees from the PANTHER database^51^, spanning 143 species across the Tree of Life. Each tree was reconciled with the corresponding species phylogeny and annotated with speciation and duplication events. As expected, branches following duplications were significantly longer than those following speciation nodes. Across trees, the mean paired difference between the per-tree average post-duplication and post-speciation branch lengths was 0.209 (Wilcoxon signed-rank test, *p* < 1 × 10^−16^) **(Supplementary Fig. 13a)**, consistent with previous studies^52–54^. To focus the enrichment analysis on trees with sufficient duplication signal and informative internal structure, we applied two quality filters: at least 10% of internal nodes had to be annotated as duplications, and the ratio of total internal to terminal branch length had to be at least 0.1. This yielded 10,660 trees for downstream analysis.

We applied PhytClust to these 10,660 trees, allowing it to automatically select the optimal non-trivial number of clusters *k*\*. PhytClust showed a strong enrichment for duplication-rooted clusters (**Fig. 6a**; Methods). Nearly 48.9% of all trees (5,212) had enrichment scores at or above the 95th percentile of their respective null distributions, and 37.9% (4,040) exceeded the 99th percentile. On average, duplication-rooted clusters comprised 32.0% of PhytClust partitions, compared with 19.8% expected under permutation of duplication labels (*p* = 0.010, one-sided permutation test; **Fig. 6b**). Although only 25.8% of trees (2,751) were individually significant after Benjamini–Hochberg correction^55^, reflecting limited per-tree power, the consistent right-shift across trees **(Fig. 6a)** indicates a genome-wide bias toward duplication-rooted clustering.

**Fig 6:**
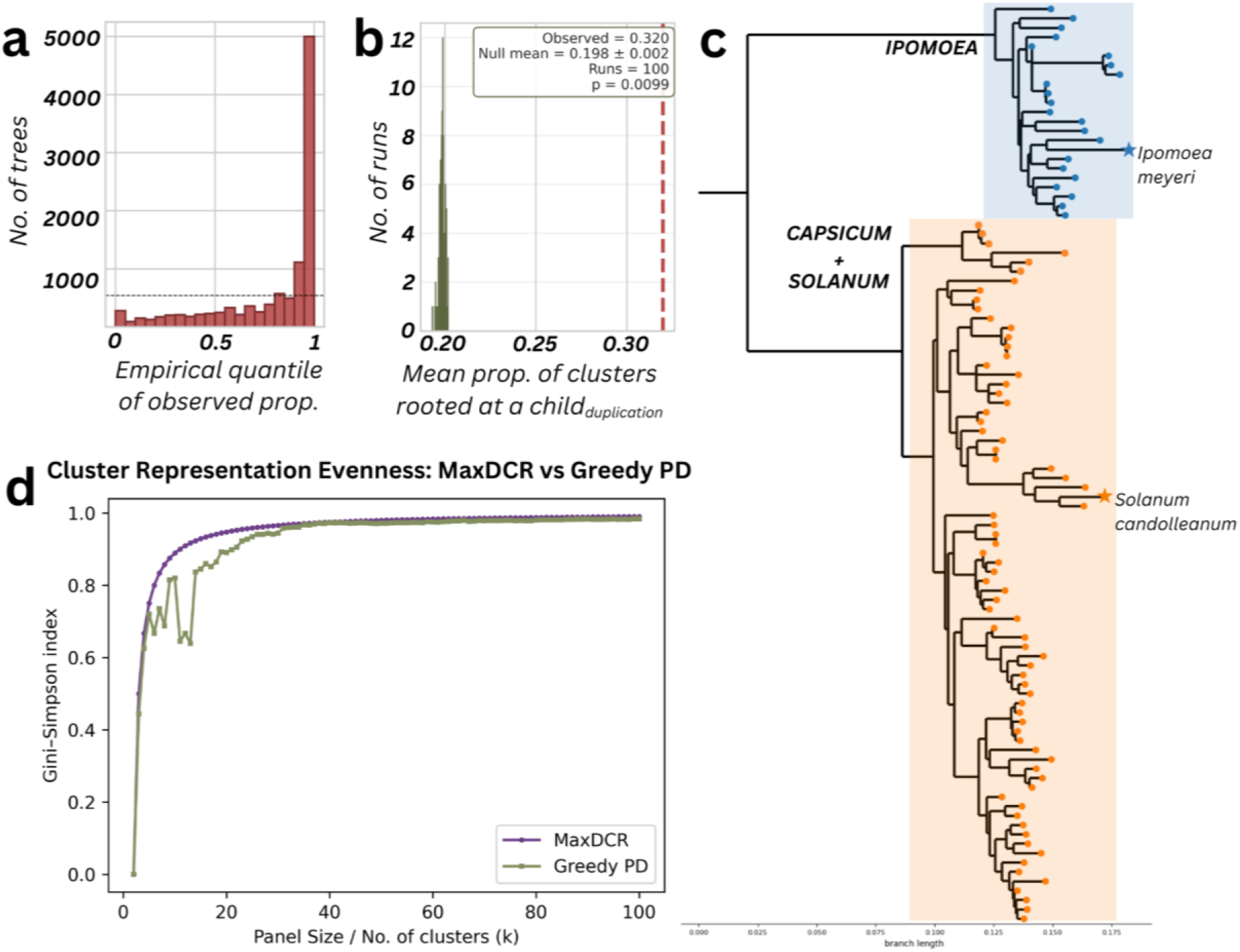
(a) Enrichment of duplication-rooted clusters across trees. Histogram showing, for each tree, the empirical quantile of the observed proportion of PhytClust clusters whose MRCA’s parent node is annotated as a duplication. (b) Global enrichment test. Null distribution (100 permutations) of the mean across trees of the proportion of duplication-rooted clusters, with the observed mean indicated by a vertical line. (c) Example section of a phylogenetic tree inferred from 185 species of crop wild relatives of Colombia, highlighting three large clusters (Solanum, Capsicum, Ipomoea) identified by PhytClust. The maximally divergent cluster representatives (MaxDCR) for each cluster are annotated. (d) Comparison of representative-panel performance across panel sizes k, showing Gini– Simpson evenness of represented clusters for MaxDCR and the greedy-PD baseline.

These findings demonstrate that PhytClust can detect biologically meaningful patterns of accelerated evolution directly from tree structure, without explicit duplication annotations. This capacity highlights its utility for large-scale analyses of gene family evolution, where duplication and diversification are pervasive. In the following section, we illustrate how such cluster outputs can inform downstream functional analyses.

### Cluster-based optimisation of phylogenetic representative selection

Phylogenetic indices are widely used to quantify biodiversity and guide conservation or sampling decisions. Phylogenetic diversity (PD) is the total branch length spanned by a selected set of taxa. Maximizing PD is a common strategy for retaining evolutionary history^56–58^.

However, PD is agnostic to which taxa contribute to that history. Selecting several closely related species from a deep clade can yield high PD while overlooking evolutionarily distinct lineages. This limitation is also relevant in practical tasks such as reference panel design, where the aim is to choose representatives such that the distance from each taxon to its nearest selected representative is small.^59^.

Because PD does not explicitly enforce representation across major lineages, we implemented a cluster-based selection strategy using PhytClust. We first partition the phylogeny into clusters and then select at most one representative from each cluster, thereby preventing multiple representatives from being drawn from the same cluster. This enforces broader lineage coverage without fixing the panel size *a priori*. Within each cluster, we then select the MaxDCR (maximally divergent cluster representative)—the taxon farthest from the cluster’s MRCA—to maximize internal diversity.

Applied to a 185-species phylogeny of Colombian crop wild relatives—wild plant taxa closely related to cultivated crops—spanning 13 families in 11 clades^60^, PhytClust yielded 11 clusters (*α* = 5.0) **(Supplementary Fig. 13b)**, including *Ipomoea, Capsicum*, and *Solanum* **(Fig. 6c)**. Using MaxDCR, the resulting representative sets showed markedly improved evenness of phylogenetic coverage relative to a standard greedy-PD baseline, while maintaining near-identical total PD. Across panel sizes *k* ∈ [2,100], MaxDCR achieved higher Gini–Simpson^61^ evenness of represented clusters **(Fig. 6d)**, indicating that PD-based selection alone can concentrate representation in a small number of clades. This occurs when adding another taxon from an already represented, internally diverse clade contributes more previously uncovered branch length than adding the first representative from a shallower unrepresented clade. Meanwhile, total PD differed by only −1–2% (median) across *k*, showing that greater representational balance does not substantially reduce evolutionary breadth.

By integrating PhytClust-derived clusters with established diversity indices, we obtain cluster-aware representative panels that reduce coverage gaps, improve interpretability, and retain maximal phylogenetic diversity.

## Discussion

Splitting phylogenetic trees into optimal monophyletic partitions is a common problem in many biological domains, yet no algorithm can recover a uniquely correct answer from the tree alone: the definition of a biologically meaningful cluster depends inherently on the application, and any method necessarily encodes a modeling choice about what “good” clustering means. Rather than treating this as a limitation to be hidden, PhytClust makes its criterion explicit and intuitive. It builds on a principle shared across clustering traditions: members of the same cluster should be relatively similar to one another, while different clusters should be relatively distinguishable. For a given *k*, PhytClust identifies the monophyletic partition that minimizes within-cluster dispersion—the sum of patristic distances from each tip to its cluster’s MRCA—which for a fixed tree equivalently maximizes between-cluster separation. We chose this measure over the sum of unique branch lengths within each cluster because the former weights each branch by the number of descendant tips it subtends and therefore penalizes large but internally diffuse clades more strongly. Because the tip-to-MRCA sum grows with the number of tips in a cluster, the objective implicitly favors more balanced partitions, which may explain, for example, the subdivision of the larger *Lokiarchaeota* into three groups while smaller phyla are merged. The goal is not to impose a single correct answer but to provide a standardized, reproducible starting point that reflects broadly shared intuitions about what a coherent group looks like—one that can then be refined in light of domain knowledge.

Compared with the threshold-dependent and embedding-based methods surveyed in the Introduction, PhytClust operates directly on rooted tree geometry without requiring user-defined cutoffs or intermediate representations. We ran broad silhouette-based sweeps for TreeCluster and PhyloPart following prior benchmarking practice, but silhouette may not be the best fit for phylogenetic structure, and wider sweeps increase runtime, underscoring the value of a threshold-free approach. Complementary phylogeny-aware approaches^62,63^ incorporate extrinsic information such as environmental or sample metadata, whereas PhytClust clusters solely on tree topology and branch lengths. Despite these different design choices, PhytClust consistently reproduces partitions similar to those obtained by expert manual curation, suggesting that its criterion aligns well with human expectations of biologically meaningful groups. Although our empirical estimates of signal strength *α* were obtained primarily from viral phylogenies, PhytClust achieved high agreement with expert-curated classifications on the avian and Asgard archaea phylogenies, demonstrating that it can recover biologically meaningful partitions even at signal strengths well below those observed in viral datasets.

Several limitations should be noted. As with all methods on rooted trees, PhytClust’s partitions depend on the chosen rooting, since clusters are defined through monophyly and MRCAs. PhytClust can be applied to both substitution-based and time-scaled phylogenies; for the same taxa, these may yield different partitions when rate heterogeneity makes the relationship between substitutions and time non-linear. Furthermore, while the partition for each fixed *k* is computed exactly, the selected *k*\* is optimal only with respect to our combined cluster-validity score, and users whose biological question favors a different resolution can inspect the full set of optimal partitions across *k*. The multi-resolution stratification we provide supports this by surfacing well-supported partitions at several hierarchical levels. PhytClust scales quadratically with the number of leaves, and despite our efficient implementation handling trees with over 10^5^ taxa, it may not be directly applicable to metagenomics datasets with millions of taxa^64^. For such cases, collapsing clades, iterative subclustering, or capping the maximum *k* may be practical strategies. Finally, PhytClust identifies strictly monophyletic clades, which may not suit applications where lineage definitions rest on shared mutations rather than strict common descent, or where incomplete sampling and recombination violate monophyly. In such cases, methods accommodating paraphyletic groupings, such as PhyCLIP, may be better suited.

Although TreeCluster completes faster in single-threshold runs, PhytClust performs a more comprehensive optimization by simultaneously computing optimal partitions for all possible values of *k*. Most of its computational cost arises from constructing the dynamic programming table; retrieving any fixed-*k* partition adds only minimal overhead. Thus, even within comparable runtimes, PhytClust performs a substantially more exhaustive search.

Future work could improve both computational and methodological scope. A compiled implementation of the dynamic-programming core could reduce Python overhead, especially when combined with parallelization. Methodologically, it would be valuable to develop uncertainty-aware extensions using branch-support values or bootstrap/posterior tree samples, to address root uncertainty and unrooted trees, and to extend PhytClust to phylogenetic networks that can capture reticulate processes such as hybridization, introgression, and horizontal gene transfer^65^.

In summary, PhytClust provides a standardized, reproducible framework for partitioning rooted phylogenetic trees that is exact for any fixed number of clusters, scales to trees with over 100,000 taxa, and yields biologically interpretable, deterministic clusters across diverse domains without user-defined distance thresholds.

## Methods

### Phylogenetic tree: definitions and notation

Let *T* = (*V, E*) be a rooted phylogenetic tree with root *r*, where *V* is the set of nodes and *E* is the set of edges. The leaves *L*(*T*) ⊆ *V* represent the observed taxa, and *N* = ∣ *L*(*T*) ∣ is the number of leaves. Each edge *e* ∈ *E* has non-negative length *w*(*e*) ≥ 0, representing the branch length in the input tree (for example, evolutionary divergence or time). For any two nodes *x, y* ∈ *V*, let *d*(*x, y*) be the sum of edge lengths along the unique path between them. For any node *v* ∈ *V*, let *T*_*v*_ denote the subtree rooted at *v*, and let *L*(*v*) denote its leaf set. If *v* is internal, let *ch*(*v*) = {*v*_1_, …, *v*_*q*_} denote the set of child nodes of *v*. A set *C* ⊆ *L*(*T*) is an admissible cluster if either *C* = *L*(*v*) for some node *v* ∈ *V*, or, in the presence of a polytomy, *C* can be obtained by expanding that multifurcation with internal branches of length 0 and taking the leaf set of one of the resulting clades. For any non-empty set of leaves *C* ⊆ *L*(*T*), let *MRCA*(*C*) denote their most recent common ancestor in the input tree or, for clusters induced within a polytomy, the corresponding implied zero-length ancestor.

### Size of monophyletic search space

Let *n*(*v*) denote the number of monophyletic partitions of the descendant leaf set *L*(*v*) in the subtree rooted at node *v*. If *v* is a leaf, then *n*(*v*) = 1. If *v* is an internal node with children *ch*(*v*) = {*v*_1_, …, *v*_*q*_}, then

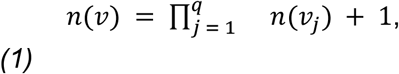

The product term counts all partitions obtained by partitioning each child subtree independently, whereas the additional + 1 accounts for the single partition in which all leaves in *ch*(*v*) = {*v*_1_, …, *v*_*q*_} are grouped together into one monophyletic cluster. For the full tree, the total number of monophyletic partitions *n*(*T*) = *n*(*r*).

To refine this count by the number of clusters, let *a*_*k*_(*v*) denote the number of monophyletic partitions of the descendant leaf set *L*(*v*) into exactly *k* clusters. For a leaf node *v*,

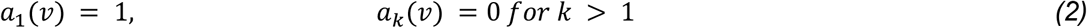

To count partitions with exactly *k* clusters, we distribute those *k* clusters across the child subtrees of *v*. If *v* has children *v*_1_, …, *v*_*q*_, let *m*_*j*_ denote the number of clusters assigned to the child subtree rooted at *v*_*j*_. Because each child subtree must contribute at least one cluster in any non-trivial partition of *L*(*v*), the values *m*_1_, …, *m*_*q*_ are positive integers satisfying 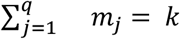. The total number of *k*-cluster monophyletic partitions of *L*(*v*) for *k* ≥ 2 is therefore

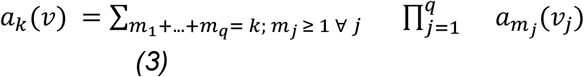

In other words, a *k*-cluster partition of *L*(*v*) is obtained by assigning a positive number of clusters to each child subtree and combining the corresponding child-subtree partitions. At the root, *a*_*k*_(*T*) = *a*_*k*_(*r*) gives the number of *k*-cluster monophyletic partitions of the full tree. In particular, *a*_1_(*T*) = *a*_*N*_(*T*) = 1.

These recurrences hold for arbitrary rooted tree topologies. To characterize the size of the search space, we next consider two extreme binary-tree topologies. In a fully imbalanced caterpillar tree with *N* leaves, each value of *k* ∈ {1, …, *N*} has exactly one monophyletic partition. Hence,

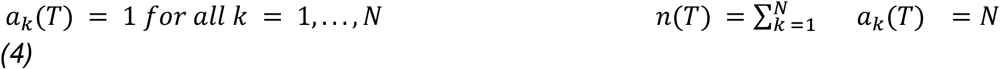

On the other hand, for a perfectly balanced binary tree of height *d*, with *N* = 2^*d*^ leaves, let *f*(*d*) denote the number of monophyletic partitions of such a tree. Then

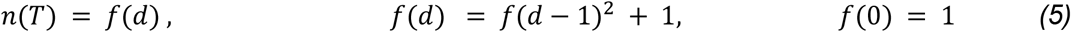

The squared term arises because the two child subtrees are themselves perfectly balanced trees of height *d* − 1, so their partitions combine independently, while the additional +1accounts for the single partition in which all leaves are grouped into one cluster. Since *f*(*d*) ≥ *f*(*d* − 1)^2^ and *f*(1) = 1, *f*(*d*) grows exponentially in the number of leaves *N* = 2^*d*^.

Thus for a rooted binary tree with *N* leaves,

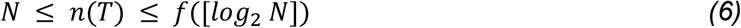

with the lower bound attained by caterpillar trees and the upper bound by perfectly balanced trees. Because perfectly balanced binary trees exist only when *N* is a power of two, the upper bound is exact at those values; for other *N*, it provides an approximate upper bound over all binary topologies. The number of monophyletic partitions therefore grows strongly with tree topology: linearly with caterpillar trees, but exponentially with the balanced trees **(Supplementary Fig. 1a)**.

### Algorithm overview

#### Finding the optimal partition for a fixed number of clusters *k*

For each value of *k* ∈ [1, *N*], we consider all monophyletic partitions of the leaf set *L*(*T*) into *k* clusters. Let *C*_*k*_ = {*c*_1_, *c*_2_, …, *c*_*k*_} denote one such partition. Our objective is to identify, among all such partitions, the one that minimizes the total within-cluster dispersion.

A valid *k*-cluster partition must satisfy three conditions:

1. each cluster *c*_*i*_ must be monophyletic;
2. the clusters consist of all leaves, so that

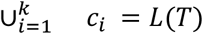

and (iii) the clusters must be mutually disjoint, so that

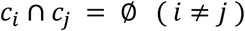

For a monophyletic cluster *c*_*i*_, let *MRCA*(*c*_*i*_) denote the root of that cluster. We define the within-cluster cost as the sum of the distances from each leaf in the cluster to its MRCA:

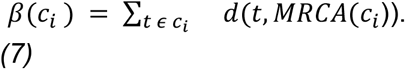

We use the sum of tip-to-MRCA distances as our measure of within-cluster dispersion because it gives a rooted-tree analogue of cluster compactness, with the MRCA acting as a cluster-specific centroid. This differs from summing each branch within a cluster only once, which measures the total size of the subtree but does not reflect how many taxa are separated by a given branch and therefore penalizes large, spread-out clades less strongly. It also differs from using all pairwise patristic distances, which is computationally less convenient for our dynamic-programming recursion because it introduces pairwise terms between taxa from different child subtrees. We chose the tip-to-MRCA formulation because it provides a good balance between biological interpretability and computational convenience, and because it aligns directly with the within-cluster term of our tree-adapted Calinski–Harabasz criterion.

The total cost of the partition *C*_*k*_ is

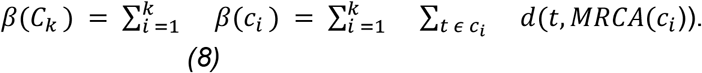

The optimal *k*-cluster partition is therefore

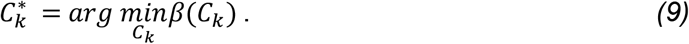

To compute 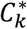 efficiently, we employ dynamic programming over the tree. Let *DP*[*v, m*] denote the minimum cost of partitioning the descendant leaf set *L*(*v*) of subtree *T*_*v*_ into exactly *m* monophyletic clusters. The DP-table is filled in post-order, so that child subproblems are solved before their parents. At the root, *DP*[*r, k*], yields the minimum cost 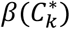 and back-pointers recover the corresponding partition.

For a leaf node *v*:

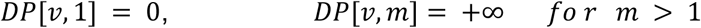

Now let *v* be an internal node with children *ch*(*v*) = {*v*_1_, …, *v*_*q*_}. If all descendants leaves of *v* are assigned to a single cluster, then the cost is simply the sum of distances from those leaves to *v:*

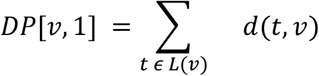

For 2 ≤ *m* ≤ |*L*(*v*)|, the optimal cost is obtained by distributing the *m* clusters across the child subtrees

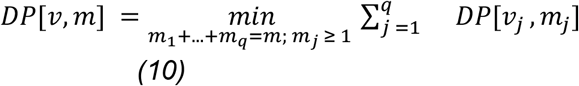

Finally,

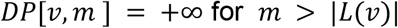

This recurrence reflects the fact that if *m* ≥ 2, then the partition of *L*(*v*) must be formed by combining partitions from its child subtrees, with each child contributing at least one cluster. For trees without polytomies, Eq. (10) gives the full recurrence. In the presence of polytomies, PhytClust extends this recursion by additionally considering clusters induced by zero-length expansions of multifurcating nodes, as described below.

An example of the DP table calculation is shown in **(Fig. 2a)**. Tree parsing and traversal were implemented using the *Bio*.*Phylo* module from Biopython^66^ and pseudocode for the dynamic programming and backtracking is provided in the Supplementary Material.

#### Selecting an optimal number of clusters using a cluster-validity index

After computing the optimal partition 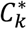 for each *k* ∈ {1, …, *N*}, we next select the most informative non-trivial number of clusters, *k\**. Because the dynamic programming objective 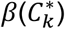 minimizes the total within-cluster dispersion, and because the total distance from all leaves to the root is fixed for a given tree, minimizing the within-cluster term is equivalent to maximizing the corresponding between-cluster term **(Supplementary Proof A)**.

To evaluate candidate resolutions, we use a tree-adapted version of the Calinski–Harabasz (CH) index^25^. In this setting, we treat the most recent common ancestor (MRCA) of each cluster as its cluster centroid and the tree root *r* as the global centroid. Because the square root of patristic branch lengths permits an embedding of leaf nodes in Euclidean space ^67^, we use branch lengths directly in place of squared Euclidean distances, replacing sums of squared Euclidean norms by sums of patristic distances on the tree. Following the usual CH index, we define the within-cluster term as

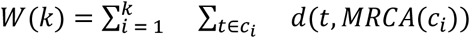

where 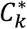 is the optimal *k*-cluster partition. We further define the total sum of distances to the root as

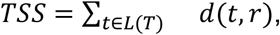

Note that 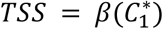, since the single-cluster partition has MRCA at the root. The between-cluster term is then

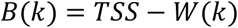

By construction, the dynamic programming objective equals the within-cluster term, so

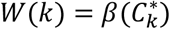

The tree-adapted Calinski–Harabasz index is therefore

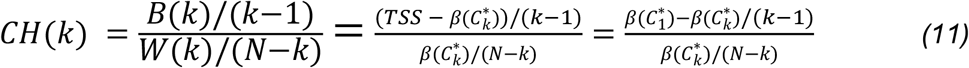

As in the standard Calinski–Harabasz index, the factors *k* − 1 and *N* − *k* normalize the between- and within-cluster terms by their respective degrees of freedom, helping to penalize overly fine partitions unless additional clusters yield sufficient separation.

As in Euclidean clustering, the CH score favors partitions with high between-cluster separation and low within-cluster dispersion. However, *CH*(*k*) may continue to increase with *k* when *W*(*k*) decreases rapidly, which can bias selection toward overly fine-grained partitions. To reduce this tendency, we combine *CH*(*k*) with the Elbow Index (EL1)^26^, which is likewise defined directly on the tree:

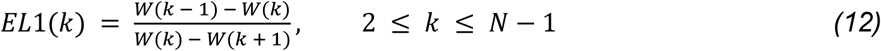

We then define a combined score

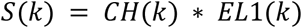

As *k* increases, *β*(*k*) monotonically decreases. The CH term favors partitions with low within-cluster cost relative to between-cluster separation, whereas EL1 highlights points at which the decrease in *β*(*k*) begins to level off. We therefore use their product *S*(*k*) to favor values of *k* that combine strong separation with diminishing returns in further cluster subdivision.

Rather than selecting the global maximum of *S*(*k*) directly, we first identify all local maxima of *S*(*k*) over the non-trivial range *k* ∈ {2, …, *N* − 1}. Each local maximum is treated as a candidate resolution. For each such peak, we consider both its height *S*(*k*) and its prominence *P*(*k*), where prominence is defined as the vertical distance between the peak and the nearest lower local minimum. Prominence therefore measures how strongly a local maximum stands out from surrounding fluctuations and helps distinguish well-supported candidate resolutions from minor local irregularities in the score profile. We then rank candidate peaks using

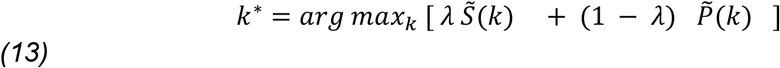

*where* 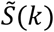 and 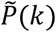 are the normalized score and prominence, respectively and *λ* ∈ [0, 1] controls the relative weight given to peak height versus peak prominence. We set *λ* = 0.7 based on a stability sweep over simulated trees **(Supplementary Fig. 14b)**; performance degraded noticeably below *λ* = 0.7, indicating that peak height relative to prominence is the more informative signal. All results reported here were using *λ* = 0.7.

PhytClust is invariant to a global positive rescaling of all branch lengths, *w*′(*e*) = *γ w*(*e*), because such a transformation multiplies all candidate partition costs by the same constant *γ*and therefore does not change either the optimal partition for fixed *k* or the selected value *k\**. By contrast, nonlinear branch-length transformations change the relative weighting of short and long branches and may therefore alter both the optimal partition and the selected clustering resolution.

#### Multiple clade levels

Each local maximum of *S*(*k*) corresponds to a candidate clustering resolution. This is useful because a single tree may support several biologically meaningful partitions at different levels of granularity, analogous to taxonomic ranks such as family, genus, and species. Rather than forcing a single optimal solution, users may therefore wish to recover a fixed number of clade levels (CLs), with one representative partition at each level.

To support this, we stratify the candidate values of *k* into *b* logarithmically spaced bins over the non-trivial range *k* ∈ {2, …, *N* − 1}, where *b* is the number of clade levels requested by the user. Specifically, we define the the interval boundaries as

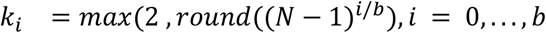

Within each interval [*k*_*i*_ − 1, *k*_*i*_], we identify the best-supported local maximum of S(k) and report the corresponding partition as the representative solution for that clade level. For example, when *N* = 1000 and *b* = 5, the resulting levels are *k* = {2, 4, 16, 63, 251, 999}.

#### Handling polytomies

For trees containing polytomies, we interpret a multifurcation as collapsed zero-length branching. Under this interpretation, clusters need not be restricted to clades rooted at nodes explicitly present in the input tree; they may also be formed by grouping subsets of the children of a polytomous node, as if these were connected by additional internal branches of length 0.

For example, consider a polytomous node with four children *A, B, C, D*. Rather than being forced to treat all four as a single cluster, PhytClust considers all possible groupings of the children— {*A, B*} and {*C, D*}, or {*B, C*} and {*A, D*}, or {*A, B, C*} and {*D*}, and so on—by implying zero-length internal branches between them. The grouping that minimizes *β* is then selected, meaning the chosen partition depends entirely on the branch-length structure of the subtrees rooted at each child. Because such implied branches do not change patristic distances, different groupings within the same polytomy can yield the same within-cluster dispersion *β*. Multiple partitions may therefore be equally optimal for a given *k*, in which case PhytClust returns one deterministic representative of the tied optima.

#### Identifying Representative Species

Phylogenetic diversity (PD) is defined as the total branch length spanned by a selected set of taxa and is widely used as a measure of evolutionary diversity. Given a PhytClust partition *C*_*k*_ = {*c*_1_, *c*_2_, …, *c*_*k*_}, we select one representative taxon from each cluster by maximizing the distance from the cluster MRCA:

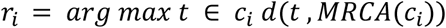

We refer to *r*_*i*_ as the maximally divergent cluster representative (MaxDCR).

### Evaluating performance using synthetic trees

#### Simulation Setup

We generated random rooted binary trees with *N* leaves using ETE3^68^, under the Proportional to Distinguishable Arrangements (PDA)^24^ distribution which is uniform over labeled rooted binary topologies. All edges were initially assigned to length 1. For a chosen value of *k*, we generated a random monophyletic target partition by recursively propagating integer labels through the tree, beginning at the root, until exactly *k* nodes were designated as target clade roots. At each step, assignments were sampled subject to feasibility constraints imposed by subtree sizes, so that no subtree was assigned more clusters than the number of leaves it contains. The descendant leaf sets of the final selected nodes defined the target clades.

We then defined signal strength *α* as the ratio of average extra-cluster to intra-cluster branch length and multiplied all extra-cluster edges by *α* ∈ [1, 50]. Thus, increasing *α* strengthens separation among the target clades while leaving within-clade branch lengths unchanged. Fixed random seeds were used for reproducibility.

As a performance metric, we used the V-Measure^69^, defined as the harmonic mean of homogeneity (*h)* and completeness (*c*) given by

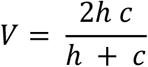

Here,

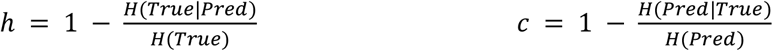

Where *H*(⋅) denotes Shannon entropy^70^ and *True* and *Pred* denote the ground-truth and predicted cluster labels. The V-measure ranges from 0 to 1, with higher values indicating better agreement between the inferred partition and the ground-truth clustering. Homogeneity is high when each inferred cluster contains leaves from only one ground-truth class, whereas completeness is high when all leaves from a given ground-truth class are assigned to the same inferred cluster. We weigh homogeneity and completeness equally.

We chose the V-Measure over the Adjusted Rand Index^71^ (ARI) because ARI is relatively insensitive to singleton clusters^69,72^ **(Supplementary Fig. 14a)**. In addition to V-Measure, we also report Normalized Mutual Information^73^ (NMI) as a supplementary validation metric (**Supplementary Fig. 14a)**.

#### Performance against random clustering

For each *k* ∈ {5,10,20} we generated 100 rooted binary trees under the PDA distribution with *N* = 100 leaves. As a monophyletic random baseline, we generated 100 random *k*-partitions per tree by sampling *k* disjoint internal nodes as described above, without using branch-length information. PhytClust was then run with a *k* fixed to the true number of clusters, and clustering performance was quantified using the V-measure for each replicate.

To provide a worst-case reference, we also used the same dynamic programming framework but reversed the optimization objective, maximizing *β*(*C*_*k*_) rather than minimizing it. This yields monophyletic partitions that are intentionally poorly separated with respect to our objective. We refer to this baseline as the *worst-separation* partition (PhytWorst). Together, the random and worst-separation baselines bracket PhytClust’s performance between unstructured random clustering and deliberately poor clustering **(Supplementary Fig. 2b)**.

#### Benchmarking against other phylogenetic clustering tools

We benchmarked PhytClust against several existing phylogeny-based clustering tools on the same simulated trees described above. We considered AutoPhy^22^, TreeCluster^16^, PhyloPart^32^, and PhyCLIP^17^. We excluded ClusterPicker, because previous studies have reported that TreeCluster and PhyCLIP generally outperform it^16,17,19^. AutoPhy is parameter-free and requires only a phylogenetic tree as input, so it was run with default settings.

PhyloPart requires a user-defined percentile threshold on global pairwise patristic distances. Following the benchmarking strategy of PhyCLIP’s study^18^, we ran PhyloPart over a grid of patristic distance thresholds from 0.05 to 0.50 in steps of 0.05. For each threshold, we obtained a clustering and computed the silhouette score on patristic distances. For each taxon *i*, the silhouette score is defined as *s*(*i*) = [(*b*(*i*) − *a* (*i*)]/*max*{*a*(*i*), *b*(*i*)}, where *a* (*i*) is the mean distance from *i* to taxa in its assigned cluster and *b* (*i*) is the smallest mean distance from *i* to taxa in any other cluster. We used the average silhouette score computed from the pairwise patristic distance matrix. Higher average silhouette scores indicate better separation between clusters relative to within-cluster cohesion. Hence, the partition with the highest silhouette score was selected for comparison.

TreeCluster offers a threshold-free mode that maximises the number of non-singleton clusters. On our simulated trees, however, this mode tended to produce clusters of size two only. To obtain more interpretable partitions, we instead used the “*sum_branch_clade*” objective and scanned thresholds from 1 to 10 in steps of 1. As for PhyloPart, we computed silhouette scores for each threshold and selected, for each tree, the partition with the highest silhouette score.

For PhyCLIP, we explored a broad parameter range: minimum cluster size fixed at 2, *γ* ∈ [0.10, 0.36] in steps of 0.05, and *FDR* ∈ {1, 2, 3, 4, 5}. For each parameter combination, PhyCLIP can return one or more candidate clusterings. We used its “*intermediate resolution*” selection mode to choose a single clustering per parameter setting, and then selected, for each tree, the solution with the highest silhouette score on patristic distances. For ease of comparison across methods, singleton leaves were treated as one-member clusters when computing external validation metrics.

For all methods, clustering performance was evaluated on each simulated tree using the V-measure, as in the main simulation study, together with the adjusted rand index (ARI) and normalized mutual information (NMI) **(Supplementary Fig. 14a)**.

To relate PhytClust’s performance to properties of the underlying tree, we computed two additional summary statistics for each simulated tree. First, we defined the backbone tree by collapsing each true target cluster to a single tip at its MRCA, thereby removing all branches below the cluster roots while preserving the higher-level topology. We quantified backbone balance using the normalized Sackin index^74^, defined as the sum of root-to-tip depths across all leaves normalized by tree size.

Second, we quantified heterogeneity in true cluster sizes using the Gini coefficient of the cluster-size distribution. Here, the input values were the numbers of leaves in each ground-truth cluster. The Gini coefficient ranges from 0, when all clusters are equal in size, to 1, when cluster sizes are maximally unequal. In **Supplementary Fig. 2d**, we relate the V-measure achieved by PhytClust to backbone imbalance and cluster-size Gini.

To summarize PhytClust’s recovery across signal strengths for a given simulated tree, we computed the area under the V-measure-versus-*α* curve (AUC) using the tested α grid. Higher AUC values indicate that PhytClust achieves higher V-measure over a wider range of signal strengths, including at lower *α*. In **Supplementary Fig. 2d**, we plot cluster-size Gini against backbone imbalance and color each point by the corresponding mean AUC, to assess how recovery varies with these two properties of the underlying tree.

### Running PhytClust on Real Trees

#### Signal strengths in real trees

To investigate whether signal strengths where PhytClust’s partitioning approaches the ground truth clusters in the simulations are also found in empirical trees, we applied PhytClust to several phylogenies. These included the Dengue (accession date: 10.04.2026), Monkeypox (accession date: 18.11.2025), Tick-borne encephalitis virus (accession date: 18.11.2025) and Tuberculosis (accession date: 10.04.2026) phylogenies from Nextstrain^29,30,33^ as well as a coronavirus (accession date: 18.11.2025) phylogeny from Dong et al.^31^. For the Dengue and coronavirus trees, PhytClust was run without a fixed *k*. For the monkeypox and tick-borne encephalitis trees, we additionally ran PhytClust with *k* constrained to match the number of named clades in the corresponding studies, enabling direct comparison with the curated nomenclature. For the tuberculosis phylogeny, PhytClust was run both without a fixed *k*, to examine the top-ranked candidate resolutions in the score profile, and with *k* fixed to 11, matching the number of major annotated lineages. For dengue, PhytClust was additionally evaluated at genotype resolution by fixing *k* = 20 to match the 20 annotated dengue genotypes. To summarize clade separation in real trees, we treated the named clades from Nextstrain, or from Dong et al. for coronaviruses, as reference clusters. For each reference clade, we identified its MRCA in the tree and defined *α* as the ratio of average extra-cluster to average within-cluster dispersion. In contrast to the simulation setting, this quantity is used here only as a descriptive summary statistic of separation in the observed tree and not as a true generative parameter. We also ran AutoPhy on these trees for comparison, however AutoPhy did not finish running within 4 hours for all phylogenies except for the tick-borne encephalitis and monkeypox phylogenies.

#### Cancer Phylogenies

For the first cancer example, we obtained the phylogenetic tree, copy-number profiles, and whole-genome-doubling (WGD) calls for patient OV-025 from McPherson et al.^41^. In the study, copy-number profiles were derived from single-cell whole-genome sequencing of ovarian cancer samples and used to reconstruct the patient’s phylogeny with MEDICC2 ^42^. Cluster WGD purity was defined as the proportion of cells within each inferred subclone that exhibited a WGD state, for example, 1xWGD and 2xWGD. We ran PhytClust to find *k*\*, using the *diploid* as an outgroup.

For each cluster, segments present in at least 85% of member cells were retained as recurrent events. To distinguish unique from shared alterations, we constructed a segment-by-cluster presence matrix by grouping identical genomic intervals across clusters. Segments present in exactly one cluster were labeled unique, whereas segments present in two or more clusters were labeled shared. Adjacent segments with identical copy-number states were merged when their intervals were contiguous, yielding a minimal non-redundant event set for each cluster. Each merged segment was then classified into standard copy-number categories, such as gain, loss, copy-neutral loss of heterozygosity, homozygous deletion, or amplification, based on total and allele-specific copy number. For functional annotation, merged segments were intersected with curated oncogene and tumour-suppressor gene coordinates using pybedtools^75^. Genes overlapping a unique segment were reported as candidate cluster-specific driver alterations.

For the second cancer example, we obtained the breast-cancer phylogeny from Minussi et al.^13^ and the corresponding cluster annotations from Kaufmann et al^42^ and ran PhytClust with *k* = 6 and *k* = 10.

#### Avian Phylogeny

The Avian tree and corresponding family-level annotations were obtained from Stiller et al.^46^. In the study, whole-genome alignments of 363 bird species representing 218 families were used to extract intergenic regions. These regions were divided into 2-kb windows, from which the most complete 1-kb segment was selected in each window. Maximum-likelihood gene trees were then inferred for individual loci and combined to construct a coalescent-based species tree, followed by post-processing.

We ran PhytClust on this tree using our clade-level procedure with *b* = 5 logarithmically spaced intervals over the range of candidate *k* values, yielding five clade levels (CL1–CL5) from coarse to fine resolution. For each level, we retained the locally optimal partition identified by our combined cluster-validity index.

To compare PhytClust clusters with the taxonomy reported by Stiller et al., we treated their higher-level groups as reference labels and compared them with the CL3 partition using the V-measure, as in the simulation study. To compare within-cluster branch-length costs, we also computed *β* for the reference taxonomy by treating each taxonomic group as a cluster, identifying its MRCA in the tree, and summing branch lengths from each member species to that MRCA, analogous to the quantity minimized by PhytClust.

#### Asgard Phylogeny

The Asgard phylogeny was obtained from Liu et al.^9^. In the study, the tree was inferred from 209 core clusters of orthologous genes (COGs) conserved across the Asgard archaea. Protein sequences corresponding to these COGs were extracted from 162 Asgard genomes, aligned using MUSCLE and the concatenated alignment was used to infer a maximum-likelihood phylogenetic tree using IQ-Tree with the LG + F + R10 model. We ran PhytClust on this tree without fixing *k*, allowing the method to infer the optimal number of clusters *k\**. To quantify agreement with the phylum-level groupings defined by Liu et al., we treated their taxonomic assignments as reference labels and computed the V-measure between these labels and the PhytClust partition.

#### Phylogeny of crop wild relatives (CWRs)

The phylogeny used for representative-species selection was obtained from González‐Orozco et al.^60^ who assembled a maximum-likelihood tree of 185 Colombian crop wild relatives, i.e. wild plant species closely related to cultivated crops, spanning 13 families and 11 major clades using seven genetic markers and RAxML^76^. We applied PhytClust to this tree and considered the global clustering solutions for all *k* from 2 to 100. For each *k*, we treated the *k* PhytClust clusters as the units to be covered and selected one representative taxon per cluster using the MaxDCR procedure implemented in PhytClust.

As a cluster-agnostic baseline, we implemented a standard greedy phylogenetic diversity (PD) algorithm. For a given panel size *q*, we first selected the tip with the longest root-to-tip path, and then iteratively added the tip that maximised the marginal gain in Faith’s PD^58^, defined as the sum of branch lengths in the minimal subtree connecting the selected taxa, until we get *q* tips. For each *k*, we computed total PD for the MaxDCR and greedy-PD panels and recorded the relative difference *(MaxDCR* − *greedy) / greedy*. These differences were summarized across *k* using the median.

To quantify how evenly each panel covered the PhytClust clusters, we computed a Gini– Simpson evenness index for each panel and value of *k*. For a given panel, we counted how many selected taxa fell into each cluster, converted these counts into frequencies *p*_*i*_ and calculated

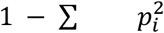

This quantity is small when most representatives come from a single cluster and increases as representation becomes more even across clusters. We compared Gini–Simpson values for MaxDCR and the greedy-PD baseline across panel sizes *k* ∈ [2, 100].

### Duplication nodes

We downloaded 15,693 reconciled gene trees from PANTHER v18.0^51^, in which each internal node is annotated as either a speciation or duplication event. To compare branch lengths following duplication and speciation events, we summarized branch lengths within each tree. Specifically, we computed the mean length of branches descending from duplication nodes and the mean length of branches descending from speciation nodes, and then used a one-sided Wilcoxon signed-rank test across trees to test whether post-duplication branches are longer than post-speciation branches.

For each tree, we also computed (i) the proportion of duplication nodes among all internal nodes and (ii) the ratio of total internal to total terminal branch length. These quantities were used only as quality filters for the downstream enrichment analysis: we retained trees in which at least 10% of internal nodes were annotated as duplications and the internal-to-terminal branch-length ratio was at least 0.1. This excluded trees with very weak duplication signals or with internal branch lengths that were small relative to terminal branches, in which PhytClust tends to return very fine partitions and cluster roots are less informative for testing enrichment. Applying these filters yielded 10,660 trees for downstream analysis.

For each retained tree, we ran PhytClust and used the top-ranked global solution *k\** as the clustering for that tree. For every cluster, we identified its MRCA and examined the parent node of that MRCA. If this parent was annotated as a duplication event (*“D=Y”* in the PANTHER annotation), we classified the cluster as duplication-rooted. For each tree we then calculated the proportion of clusters that were duplication-rooted.

To test whether duplication-rooted clusters were more frequent than expected by chance, we constructed a tree-specific permutation null model. For each tree, we held the PhytClust partition fixed, removed the duplication labels from internal nodes, and randomly reassigned the duplication label to the same number of internal nodes as in the original tree. We then recomputed the proportion of duplication-rooted clusters under this relabelling. This random reassignment was repeated 100 times per tree. The tree-wise permutation p-value was defined as the fraction of permuted proportions that were at least as large as the observed proportion, with a +1 correction applied to both numerator and denominator to avoid zero p-values. These p-values were then adjusted across trees using the Benjamini–Hochberg correction, and both raw and FDR-adjusted values were reported. For visualization, we also recorded, for each tree, the empirical quantile of the observed proportion relative to its 100 permuted values. Under the null, these quantiles are expected to be approximately uniformly distributed.

To summarize enrichment across trees, we additionally performed a global permutation test. For each random relabelling, we averaged the duplication-rooted proportion across all trees and compared the observed mean with the resulting null distribution using a right-tailed permutation test.

### Computational Performance and Technical Framework

Runtime was evaluated on randomly generated rooted binary trees with up to 100,000 leaf nodes using the ETE3 library^68^. Runtime measurements covered the complete workflow described in the benchmarking section, from reading a Newick tree to processing and saving the optimal clustering solution. Each external software package was configured as described in the simulation study. Scaling exponents were estimated by linear regression of log_10_*T* against log_10_*N* on the largest trees (*N* ≥ 5,000). We additionally used the archaeal and bacterial phylogenies from the GTDB database (release 220)^30^ to evaluate PhytClust on large empirical trees.

All benchmarks were run locally on a Lenovo ThinkPad X1 with 32 GB RAM and a 12th Gen Intel Core i7-1260P processor, to reflect typical desktop use. Larger trees can be processed with access to additional memory or high-performance computing resources.

## Data and Code availability

The source code used to perform all simulations and generate the figures presented in this manuscript is publicly available at **10.5281/zenodo.19731229**. The PhytClust software package is available through Conda or from the official Bitbucket repository at https://bitbucket.org/schwarzlab/phytclust. To facilitate interactive exploration of clustering results, PhytClust is additionally distributed with a web-based visualisation interface built on FastAPI^77^ and Uvicorn^78^, accessible from the same repository. The interface allows users to upload phylogenetic trees, inspect clustering solutions across resolutions, and export publication-quality figures without requiring programmatic access.

## Acknowledgements

This work was funded by BMBF as “SATURN3: **S**patial **A**nd **T**emporal Resol**u**tion of Intratumoural Heterogeneity in **3** hard-to-treat Ca**N**cers”, Project number 01KD2206L.

Roland F. Schwarz is a Professor at the Cancer Research Center Cologne Essen (CCCE) funded by the Ministry of Culture and Science of the State of North Rhine-Westphalia.

This work was partially funded by the German Ministry for Education and Research as BIFOLD - Berlin Institute for the Foundations of Learning and Data (ref. 01IS18025A and ref 01IS18037A). This work was supported by the Bruno and Helene Jöster Foundation as “CLONETRAC - Tracking the clonal dynamics of cancer through treatment at the single-cell level”.

We furthermore thank the ITCC (IT Center University of Cologne) for providing compute resources on the DFG-funded HPC (High Performance Computing) system RAMSES (Research Accelerator for Modeling and Simulation with Enhanced Security) as well as support (DFG funding number: INST 216/512-1 FUGG).

The authors acknowledge support from the Helmholtz Association. We would also like to thank the authors that contributed data to Nextstrain to allow us to use well-reconstructed phylogenies.

## Author contributions

K.G., T.L.K., and R.F.S. conceived and designed the study. E.B. contributed to the initial project design. K.G. and R.F.S. developed the method. K.G. implemented the software, carried out the simulations and empirical analyses, and generated the figures. M.C.C. and C.B.S. contributed mathematical input to the manuscript. C.D. and A.M.A. provided data and contributed to the development, interpretation, and writing of the corresponding section. K.G., T.L.K., and R.F.S. interpreted the results and revised and edited the manuscript. R.F.S. supervised the study. All authors read and approved the final manuscript.

## Notes

https://zenodo.org/records/19731229

https://github.com/ICCB-Cologne/PhytClust

### Competing Interest Statement

The authors have declared no competing interest.

### Summary of Updates

updated PhytClust's Repository URL and Zenodo link Restructured Results and Methods section

## References

1. Wiley, E. O. Why Trees Are Important. Evol. Educ. Outreach 3, 499–505 (2010).

2. Kapli, P., Yang, Z. & Telford, M. J. Phylogenetic tree building in the genomic age. Nat. Rev Genet. 21, 428–444 (2020).

3. Soltis, D. E. & Soltis, P.S. The Role of Phylogenetics in Comparative Genetics. Plant Physiol. 132, 1790–1800 (2003).

4. Scossa, F. & Fernie, A. R. Ancestral sequence reconstruction - An underused approach to understand the evolution of gene function in plants? Comput. Struct. Biotechnol. J. 19, 1579–1594 (2021).

5. Ng, J. & Smith, S. D. How traits shape trees: new approaches for detecting character state‐dependent lineage diversification. J. Evol. Biol. 27, 2035–2045 (2014).

6. Beerenwinkel, N., Schwarz, R. F., Gerstung, M. & Markowetz, F. Cancer Evolution: Mathematical Models and Computational Inference. Syst. Biol. 64, e1–e25 (2015).

7. Li, T. et al. Phylogenetic supertree reveals detailed evolution of SARS-CoV-2. Sci. Rep. 10, 22366 (2020).

8. Purvis, A. & Agapow, P.-M. Phylogeny Imbalance: Taxonomic Level Matters. Syst. Biol. 51, 844–854 (2002).

9. Liu, Y. et al. Expanded diversity of Asgard archaea and their relationships with eukaryotes. Nature 593, 553–557 (2021).

10. Sun, J. et al. Recoding of stop codons expands the metabolic potential of two novel Asgardarchaeota lineages. ISME Commun. 1, 30 (2021).

11. Xie, R. et al. Expanding Asgard members in the domain of Archaea sheds new light on the origin of eukaryotes. Sci. China Life Sci. 65, 818–829 (2022).

12. Zeng, L., Warren, J. L. & Zhao, H. Phylogeny-based tumor subclone identification using a Bayesian feature allocation model. Ann. Appl. Stat. 13, 1212–1241 (2019).

13. Minussi, D. C. et al. Breast tumours maintain a reservoir of subclonal diversity during expansion. Nature 592, 302–308 (2021).

14. Tarabichi, M. et al. A pan-cancer landscape of somatic mutations in non-unique regions of the human genome. Nat. Biotechnol. 39, 1589–1596 (2021).

15. Schloss, P. D. & Westcott, S. L. Assessing and improving methods used in operational taxonomic unit-based approaches for 16S rRNA gene sequence analysis. Appl. Environ. Microbiol. 77, 3219–3226 (2011).

16. Balaban, M., Moshiri, N., Mai, U., Jia, X. & Mirarab, S. TreeCluster: Clustering biological sequences using phylogenetic trees. PloS One 14, e0221068 (2019).

17. Han, A. X., Parker, E., Scholer, F., Maurer-Stroh, S. & Russell, C. A. Phylogenetic Clustering by Linear Integer Programming (PhyCLIP). Mol. Biol. Evol. 36, 1580–1595 (2019).

18. Prosperi, M. C. F. et al. A novel methodology for large-scale phylogeny partition. Nat. Commun. 2, 321 (2011).

19. Ragonnet-Cronin, M. et al. Automated analysis of phylogenetic clusters. BMC Bioinformatics 14, 317 (2013).

20. Ortiz-Velez, A. N., Sukumaran, J., Rouzbehani, R. & Kelley, S. T. AutoPhy: Automated phylogenetic identification of novel protein subfamilies. PLOS ONE 19, e0291801 (2024).

21. McInnes, L., Healy, J., Saul, N. & Großberger, L. UMAP: Uniform Manifold Approximation and Projection. J. Open Source Softw. 3, 861 (2018).

22. Reynolds, D. Gaussian Mixture Models. in Encyclopedia of Biometrics 659–663 (Springer, Boston, MA, 2009). doi:10.1007/978-0-387-73003-5_196.

23. Neath, A. A. & Cavanaugh, J. E. The Bayesian information criterion: background, derivation, and applications. WIREs Comput Stat 4, 199–203 (2012).

24. Blum, M. G. B. & François, O. Which Random Processes Describe the Tree of Life? A Large-Scale Study of Phylogenetic Tree Imbalance. Syst. Biol. 55, 685–691 (2006).

25. Calinski, T. & Harabasz, J. A dendrite method for cluster analysis. Commun. Stat. - Theory Methods 3, 1–27 (1974).

26. Rykov, A., De Amorim, R. C., Makarenkov, V. & Mirkin, B. Inertia-Based Indices to Determine the Number of Clusters in K-Means: An Experimental Evaluation. IEEE Access 12, 11761–11773 (2024).

27. Rosenberg, A. & Hirschberg, J. V-Measure: A Conditional Entropy-Based External Cluster Evaluation Measure. in Proceedings of the 2007 Joint Conference on Empirical Methods in Natural Language Processing and Computational Natural Language Learning (EMNLP-CoNLL) (ed. Eisner, J.) 410–420 (Association for Computational Linguistics, Prague, Czech Republic, 2007).

28. Rousseeuw, P. J. Silhouettes: A graphical aid to the interpretation and validation of cluster analysis. J. Comput. Appl. Math. 20, 53–65 (1987).

29. Farris, F. A. The Gini Index and Measures of Inequality. Am. Math. Mon. 117, 851–864 (2010).

30. Parks, D. H. et al. GTDB: an ongoing census of bacterial and archaeal diversity through a phylogenetically consistent, rank normalized and complete genome-based taxonomy. Nucleic Acids Res. 50, D785–D794 (2022).

31. Hadfield, J. et al. Nextstrain: real-time tracking of pathogen evolution. Bioinformatics 34, 4121–4123 (2018).

32. Sagulenko, P., Puller, V. & Neher, R. A. TreeTime: Maximum-likelihood phylodynamic analysis. Virus Evol. 4, vex042 (2018).

33. Kutschera, L. S. & Wolfinger, M. T. Evolutionary traits of Tick-borne encephalitis virus: Pervasive non-coding RNA structure conservation and molecular epidemiology. Virus Evol. 8, veac051 (2022).

34. Dong, R., Pei, S., Yin, C., He, R. L. & Yau, S. S.-T. Analysis of the Hosts and Transmission Paths of SARS-CoV-2 in the COVID-19 Outbreak. Genes 11, 637 (2020).

35. Duffy, S. Why are RNA virus mutation rates so damn high? PLOS Biol. 16, e3000003 (2018).

36. Duchêne, S. et al. Genome-scale rates of evolutionary change in bacteria. Microb. Genomics 2, e000094 (2016).

37. Menardo, F., Duchêne, S., Brites, D. & Gagneux, S. The molecular clock of Mycobacterium tuberculosis. PLOS Pathog. 15, e1008067 (2019).

38. Vasilakis, N. et al. Genetic and phenotypic characterization of sylvatic dengue virus type 2 strains. Virology 377, 296–307 (2008).

39. Phadungsombat, J., Nakayama, E. E. & Shioda, T. Unraveling Dengue Virus Diversity in Asia: An Epidemiological Study through Genetic Sequences and Phylogenetic Analysis. Viruses 16, (2024).

40. Bielski, C. M. et al. Genome doubling shapes the evolution and prognosis of advanced cancers. Nat. Genet. 50, 1189–1195 (2018).

41. McPherson, A. et al. Ongoing genome doubling shapes evolvability and immunity in ovarian cancer. Nature 1–10 (2025) doi:10.1038/s41586-025-09240-3.

42. Kaufmann, T. L. et al. MEDICC2: whole-genome doubling aware copy-number phylogenies for cancer evolution. Genome Biol. 23, 241 (2022).

43. Feng, Y. et al. Serial innovations by Asgard archaea shaped the DNA replication machinery of the early eukaryotic ancestor. Nat. Ecol. Evol. 1–13 (2025) doi:10.1038/s41559-025-02882-6.

44. Tamarit, D. et al. Description of Asgardarchaeum abyssi gen. nov. spec. nov., a novel species within the class Asgardarchaeia and phylum Asgardarchaeota in accordance with the SeqCode. Syst. Appl. Microbiol. 47, 126525 (2024).

45. Zhou, Z., Liu, Y., Anantharaman, K. & Li, M. The expanding Asgard archaea invoke novel insights into Tree of Life and eukaryogenesis. mLife 1, 374–381 (2022).

46. Stiller, J. et al. Complexity of avian evolution revealed by family-level genomes. Nature 629, 851–860 (2024).

47. Kratter, A. W. The Howard and Moore Complete Checklist of the Birds of the World. The Auk 122, 712–714 (2005).

48. Sangster, G. et al. Phylogenetic definitions for 25 higher-level clade names of birds. Avian Res. 13, 100027 (2022).

49. Ohno, S. Evolution by Gene Duplication. (Springer, Berlin, Heidelberg, 1970). doi:10.1007/978-3-642-86659-3.

50. Innan, H. & Kondrashov, F. The evolution of gene duplications: classifying and distinguishing between models. Nat. Rev. Genet. 11, 97–108 (2010).

51. Thomas, P. D. et al. PANTHER: Making genome-scale phylogenetics accessible to all. Protein Sci. 31, 8–22 (2022).

52. Huminiecki, L. & Wolfe, K. H. Divergence of Spatial Gene Expression Profiles Following Species-Specific Gene Duplications in Human and Mouse. Genome Res. 14, 1870–1879 (2004).

53. Cusack, B. P. & Wolfe, K. H. Not Born Equal: Increased Rate Asymmetry in Relocated and Retrotransposed Rodent Gene Duplicates. Mol. Biol. Evol. 24, 679–686 (2007).

54. Pegueroles, C., Laurie, S. & Albà, M. M. Accelerated Evolution after Gene Duplication: A Time-Dependent Process Affecting Just One Copy. Mol. Biol. Evol. 30, 1830–1842 (2013).

55. Benjamini, Y. & Hochberg, Y. Controlling the False Discovery Rate: A Practical and Powerful Approach to Multiple Testing. J. R. Stat. Soc. Ser. B Methodol. 57, 289–300 (1995).

56. Owen, N. R., Gumbs, R., Gray, C. L. & Faith, D. P. Global conservation of phylogenetic diversity captures more than just functional diversity. Nat. Commun. 10, 859 (2019).

57. Grover, S., Markin, A., Anderson, T. K. & Eulenstein, O. Phylogenetic diversity statistics for all clades in a phylogeny. Bioinformatics 39, i177–i184 (2023).

58. Faith, D. P. Conservation evaluation and phylogenetic diversity. Biol. Conserv. 61, 1–10 (1992).

59. Matsen, F. A., Gallagher, A. & McCoy, C. O. Minimizing the Average Distance to a Closest Leaf in a Phylogenetic Tree. Syst. Biol. 62, 824–836 (2013).

60. González‐Orozco, C. E., Sosa, C. C., Thornhill, A. H. & Laffan, S. W. Phylogenetic diversity and conservation of crop wild relatives in Colombia. Evol. Appl. 14, 2603–2617 (2021).

61. Simpson, E. H. Measurement of Diversity. Nature 163, 688–688 (1949).

62. Washburne, A. D. et al. Phylofactorization: a graph partitioning algorithm to identify phylogenetic scales of ecological data. Ecol. Monogr. 89, e01353 (2019).

63. Czech, L. & Stamatakis, A. Scalable methods for analyzing and visualizing phylogenetic placement of metagenomic samples. PLOS ONE 14, e0217050 (2019).

64. Piñeiro, C. & Pichel, J. C. Efficient phylogenetic tree inference for massive taxonomic datasets: harnessing the power of a server to analyze 1 million taxa. GigaScience 13, giae055 (2024).

65. Huson, D. H. & Bryant, D. Application of Phylogenetic Networks in Evolutionary Studies. Mol. Biol. Evol. 23, 254–267 (2006).

66. Cock, P. J. A. et al. Biopython: freely available Python tools for computational molecular biology and bioinformatics. Bioinformatics 25, 1422–1423 (2009).

67. de Vienne, D. M., Aguileta, G. & Ollier, S. Euclidean Nature of Phylogenetic Distance Matrices. Syst. Biol. 60, 826–832 (2011).

68. Huerta-Cepas, J., Serra, F. & Bork, P. ETE 3: Reconstruction, Analysis, and Visualization of Phylogenomic Data. Mol. Biol. Evol. 33, 1635–1638 (2016).

69. Rosenberg, A. & Hirschberg, J. V-Measure: A Conditional Entropy-Based External Cluster Evaluation Measure. in Proceedings of the 2007 Joint Conference on Empirical Methods in Natural Language Processing and Computational Natural Language Learning (EMNLP-CoNLL) (ed. Eisner, J.) 410–420 (Association for Computational Linguistics, Prague, Czech Republic, 2007).

70. Shannon, C. E. A Mathematical Theory of Communication. Hubert, L. & Arabie, P. Comparing partitions. J. Classif. 2, 193–218 (1985).

71. Romano, S., Vinh, N. X., Bailey, J. & Verspoor, K. Adjusting for chance clustering comparison measures. J Mach Learn Res 17, 4635–4666 (2016).

72. Kvålseth, T. O. On Normalized Mutual Information: Measure Derivations and Properties. Entropy 19, 631 (2017).

73. M. Coronado, T., Mir, A., Rosselló, F. & Rotger, L. On Sackin’s original proposal: the variance of the leaves’ depths as a phylogenetic balance index. BMC Bioinformatics 21, 154 (2020).

74. Dale, R. K., Pedersen, B. S. & Quinlan, A. R. Pybedtools: a flexible Python library for manipulating genomic datasets and annotations. Bioinformatics 27, 3423–3424 (2011).

75. Stamatakis, A. RAxML version 8: a tool for phylogenetic analysis and post-analysis of large phylogenies. Bioinformatics 30, 1312–1313 (2014).

76. fastapi/CITATION.cff at master ·fastapi/fastapi. GitHub https://github.com/fastapi/fastapi/blob/master/CITATION.cff.

77. uvicorn/CITATION.cff at main ·Kludex/uvicorn. GitHub https://github.com/Kludex/uvicorn/blob/main/CITATION.cff.

