## Supplementary_Data for "PhytClust: Efficient and Optimal Monophyletic Partitioning of Rooted Phylogenetic Trees"

### Figures

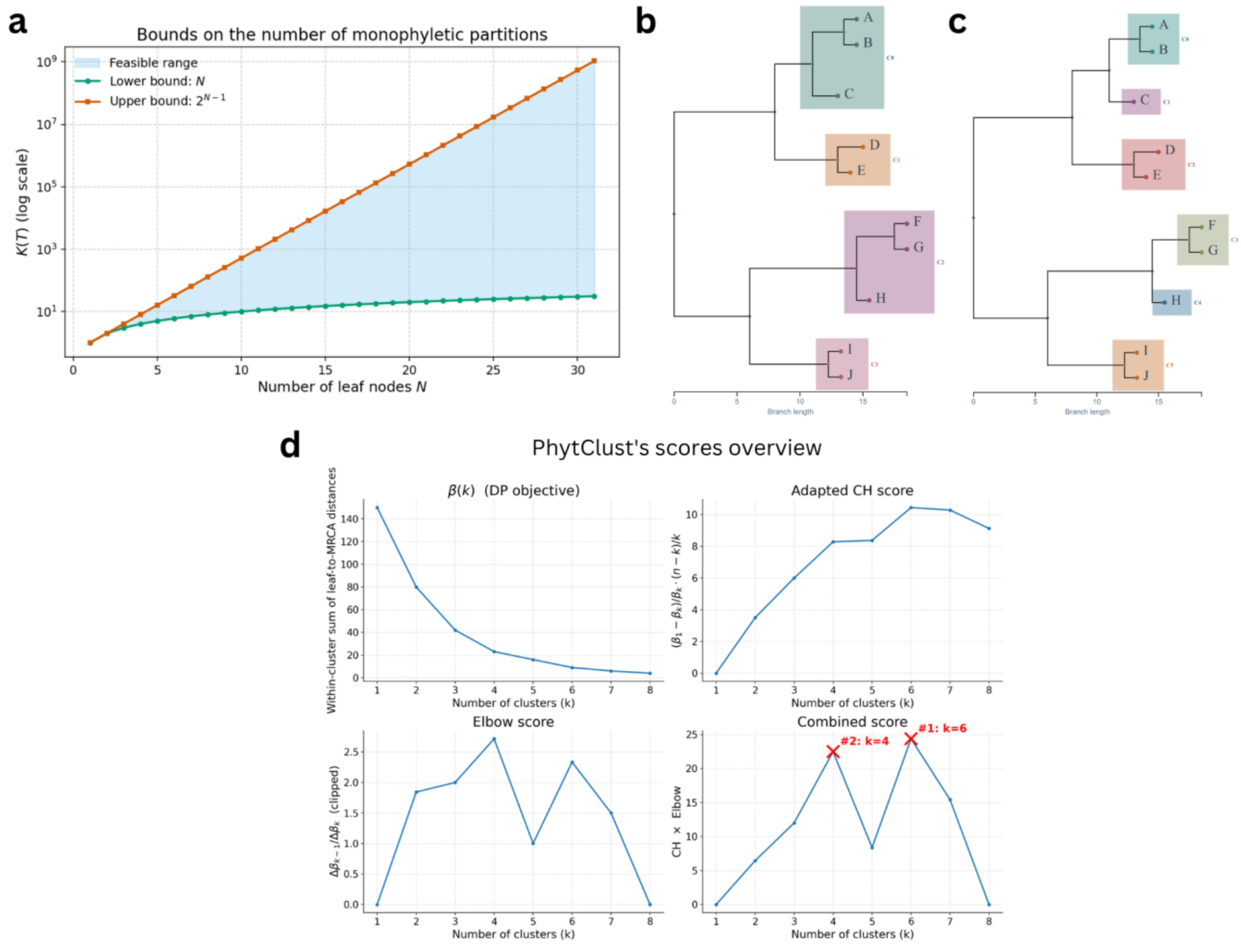


*Supplementary Figure 1: (a) Number of possible monophyletic partitions for caterpillar and perfectly balanced trees; the shaded region marks the feasible range for rooted binary trees with topologies between these extremes. (b–c) PhytClust clustering solutions for the example phylogeny shown also in panel Fig. 1a at k = 4 and k = 6. (d) Within-cluster dispersion ($\beta$), tree-adapted Calinski–Harabasz (CH), elbow (EL1), and combined cluster-quality scores for the same phylogeny, highlighting peaks at k = 4 and k = 6.*

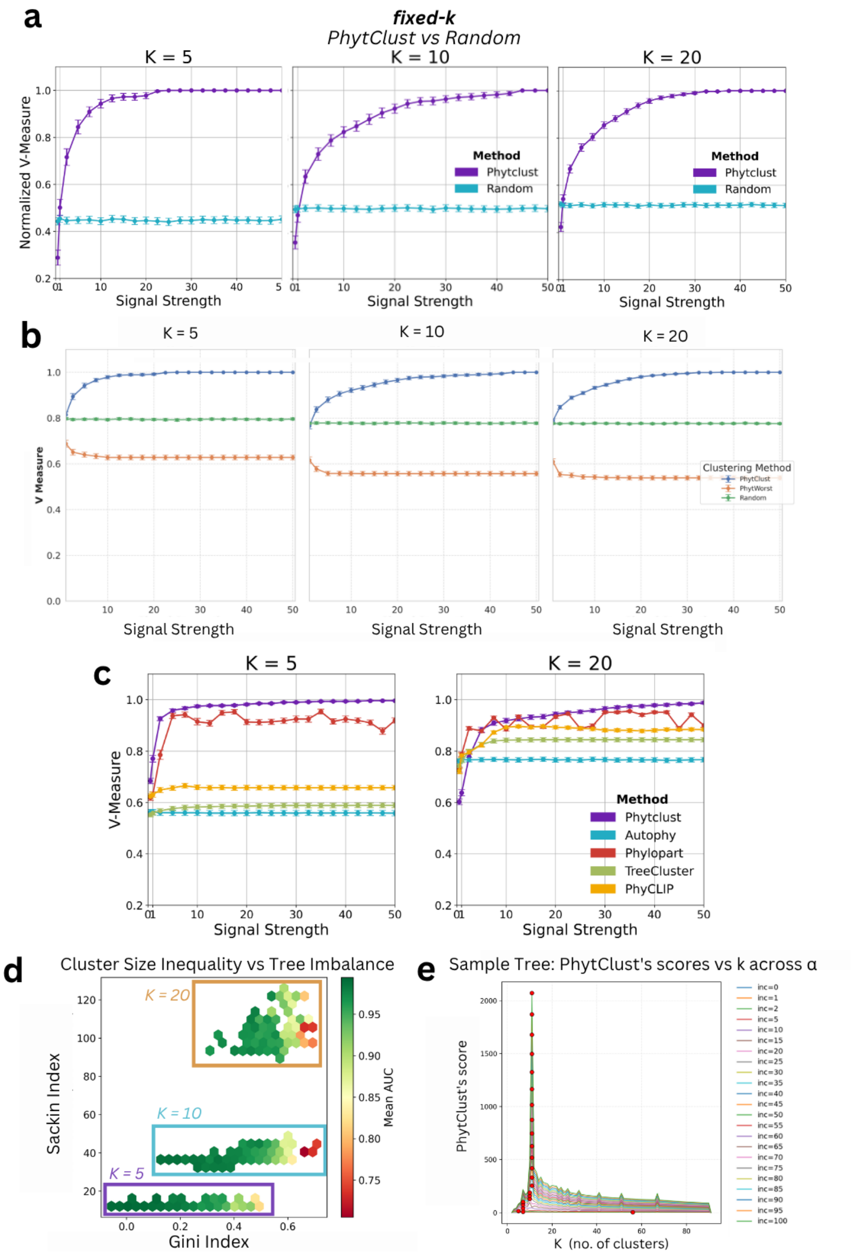


Supplementary Figure 2: (a) Simulation results comparing PhytClust with random monophyletic partitions for k = 5, k = 10, k = 20 on trees with N = 100 leaves. (b) Simulation results comparing PhytClust with random monophyletic partitions and with "PhytWorst," in which the optimization objective is reversed to maximize β(k) and intentionally produce poor clusters. (c) Simulation results for k = 5 and k = 20 on trees with N = 100 leaves across the five phylogenetic clustering tools. (d) Cluster-size inequality (Gini index) versus backbone-tree imbalance (normalized Sackin index) for simulated trees with N = 100 and k = 5, 10, 20. Points are colored by the mean AUC of the V-measure-versus-α curve for PhytClust. Higher AUC indicates better recovery across signal strengths, whereas higher Gini values indicate greater inequality in cluster sizes.(e) Sample cluster-validity index for a random tree showing the optimal k* selected by PhytClust in red. As α increases, the peak at the target k* becomes progressively more pronounced.

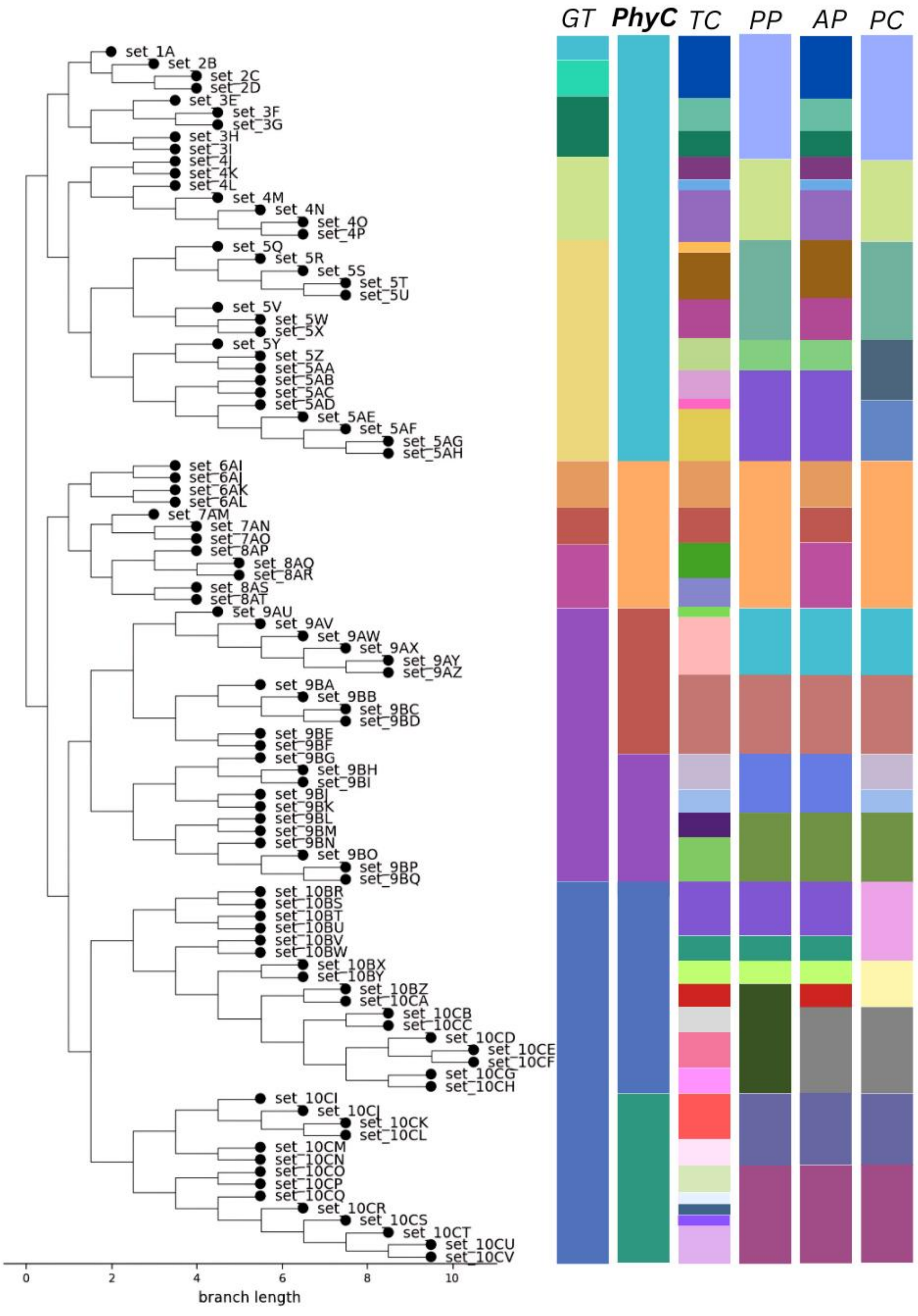


*Supplementary Figure 3: Sample tree with target clusters (GT) (N=100, k=10) in comparison with clusters identified by PhytClust (PhyC), TreeCluster (TC), PhyloPart (PP), AutoPhy (AP) and PhyCLIP (PC) when α = 1*

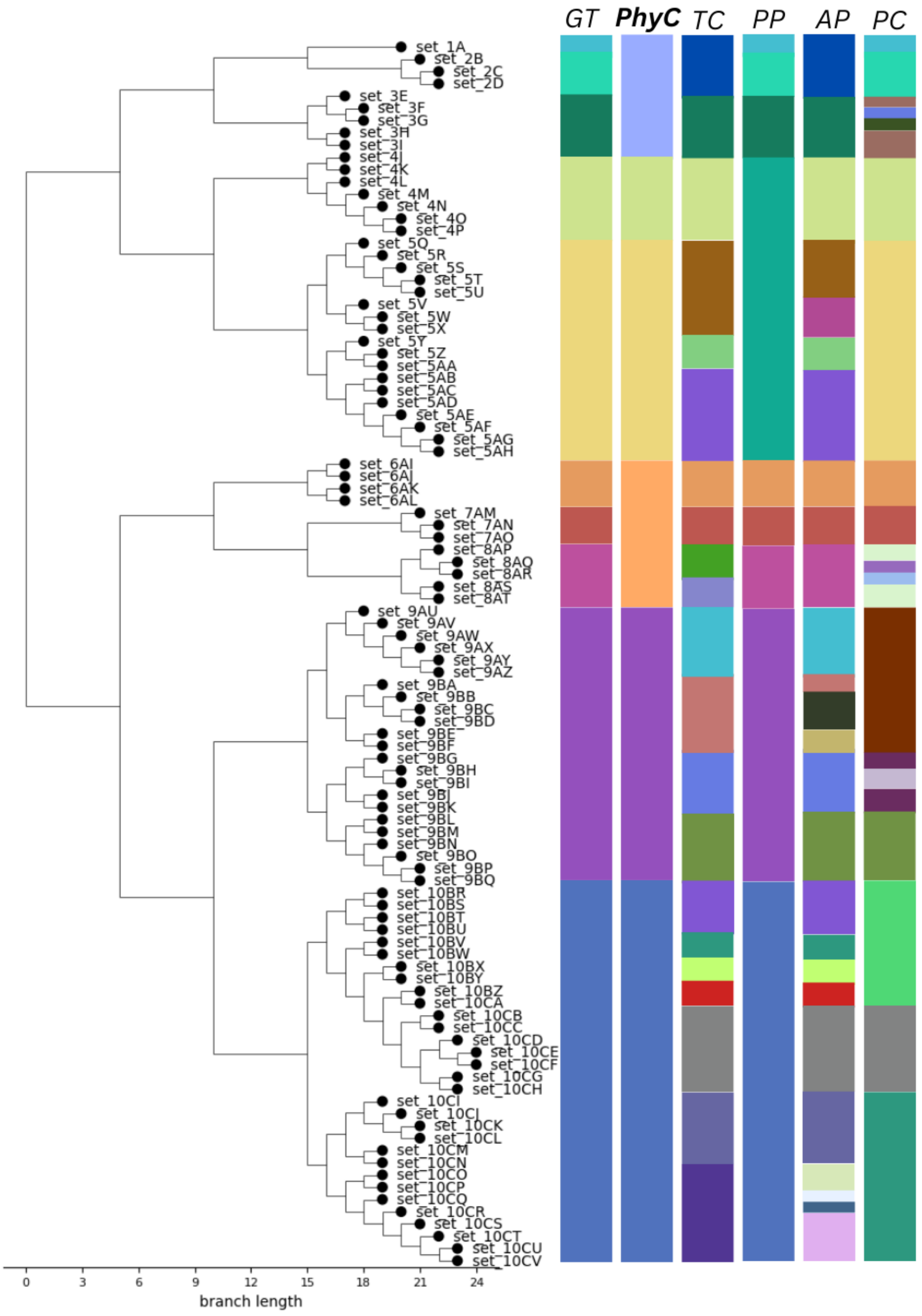


*Supplementary Figure 4: Sample tree with target clusters (GT) (N=100, k=10) in comparison with clusters identified by PhytClust (PhyC), TreeCluster (TC), PhyloPart (PP), AutoPhy (AP) and PhyCLIP (PC) when α = 5*

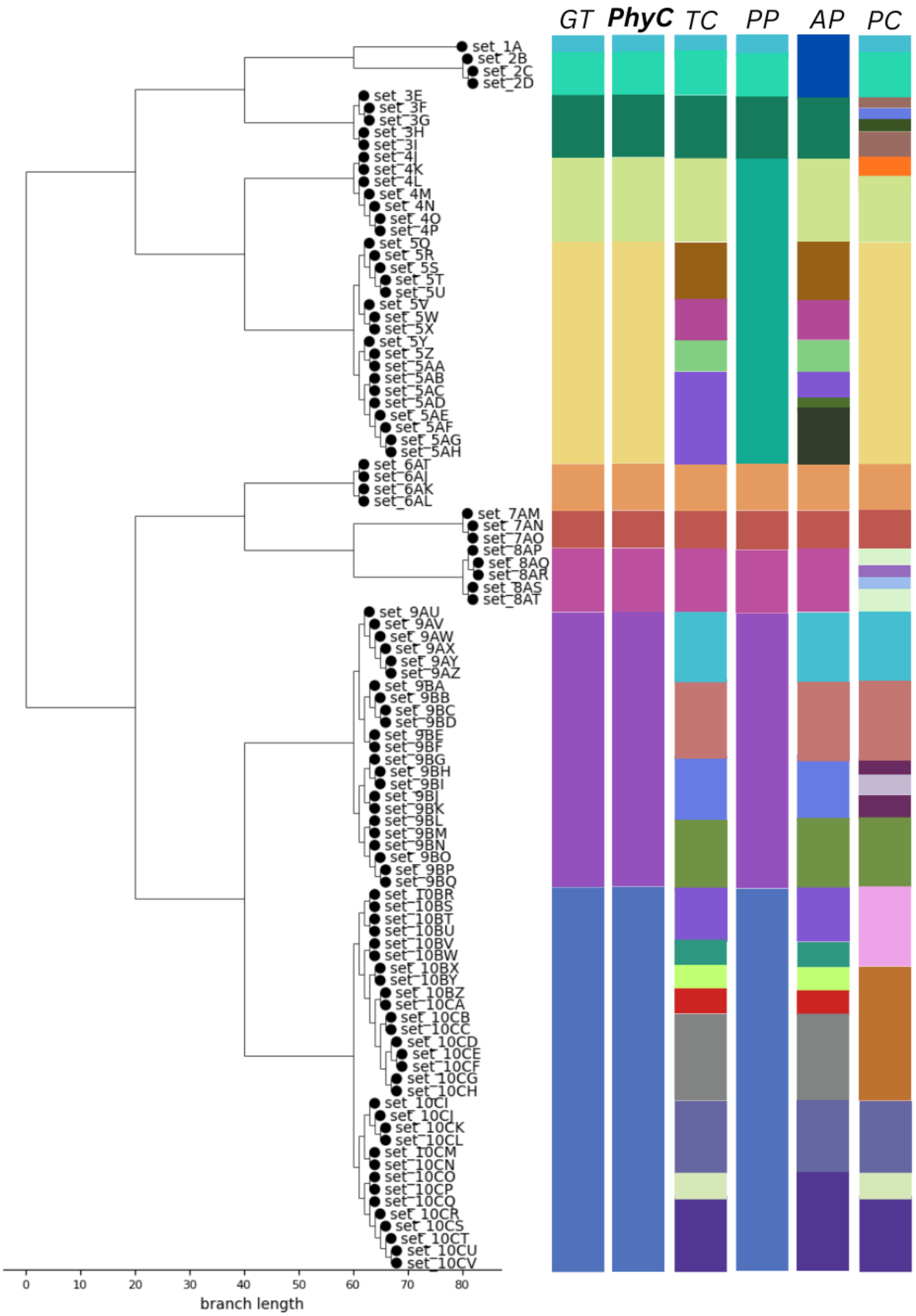


*Supplementary Figure 5: Sample tree with target clusters (GT) (N=100, k=10) in comparison with clusters identified by PhytClust (PhyC), TreeCluster (TC), PhyloPart (PP), AutoPhy (AP) and PhyCLIP (PC) when α = 10*

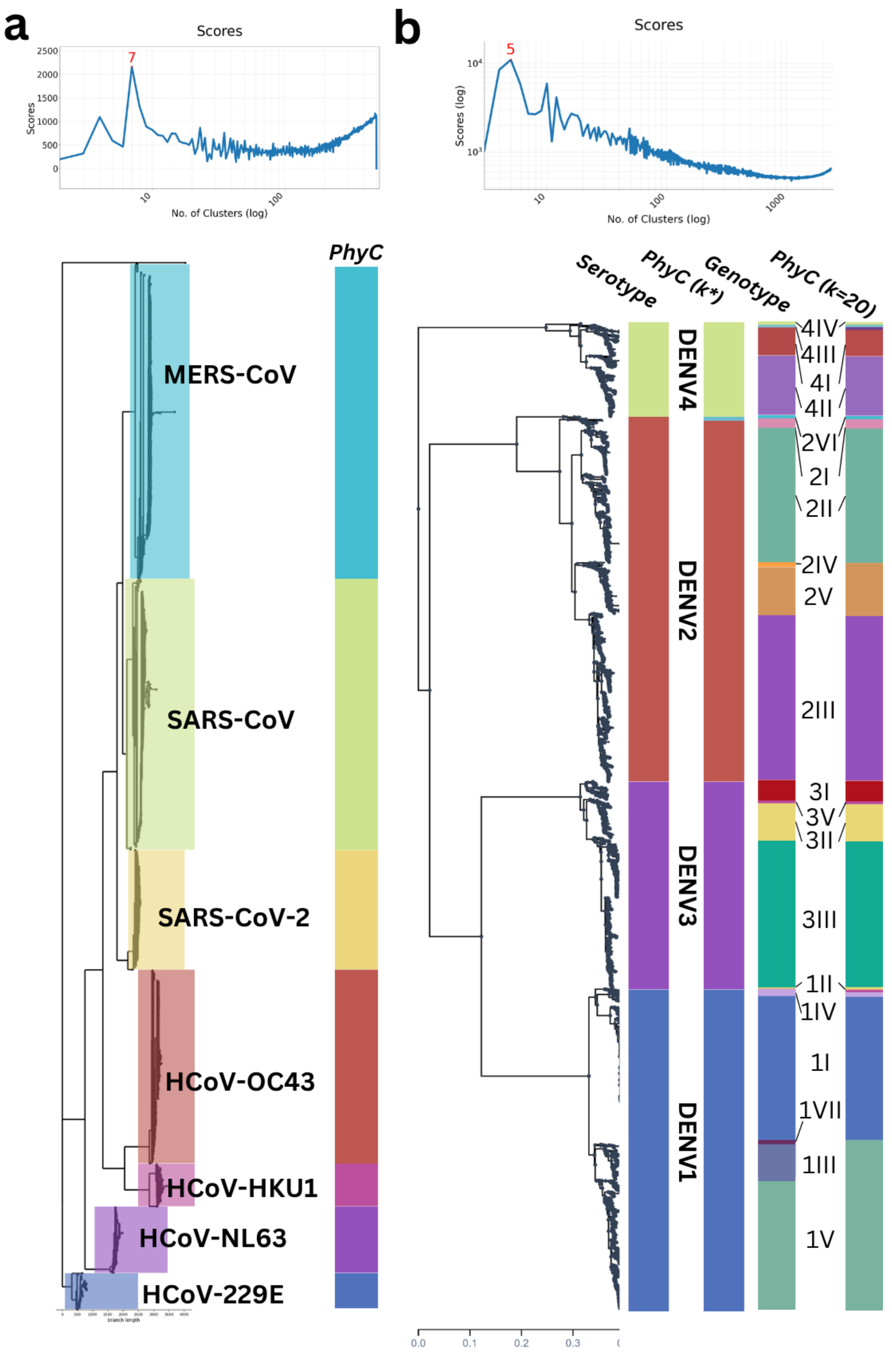


*Supplementary Figure 6: a) Coronavirus phylogeny with annotated families as well as PhytClust's chosen solution (k* = 7) and PhytClust's scores; (b) Dengue phylogeny evaluated at coarse and fine resolution. At the serotype level, the annotated Nextstrain clades are compared with PhytClust's automatically selected solution (k*= 5); the additional fifth cluster corresponds to the sylvatic DENV2 clade, yielding near-complete agreement with the serotype annotation (V-measure = 0.99). At the genotype level, the 20 annotated dengue genotypes are compared with the PhytClust partition obtained when fixing k = 20, again showing strong concordance (V-measure = 0.95). The PhytClust score profile is shown above the phylogeny.*

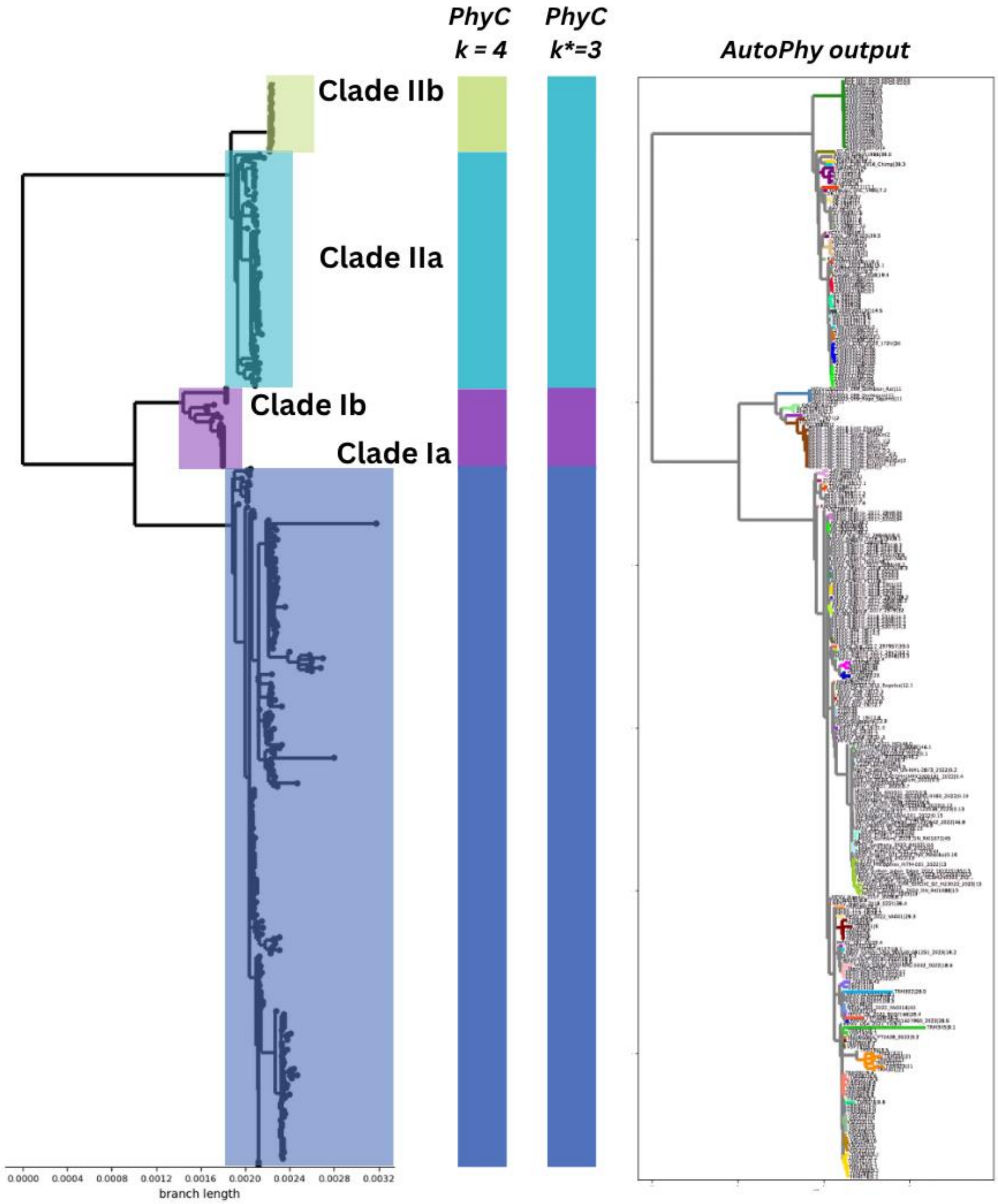


*Supplementary Figure 7: Monkeypox phylogeny with annotated clades as well as PhytClust's chosen solution (k* = 3) and PhytClust's solution for k = 4 (left), and AutoPhy's output on the same tree (right). The AutoPhy visualization is reproduced directly from the software output.*

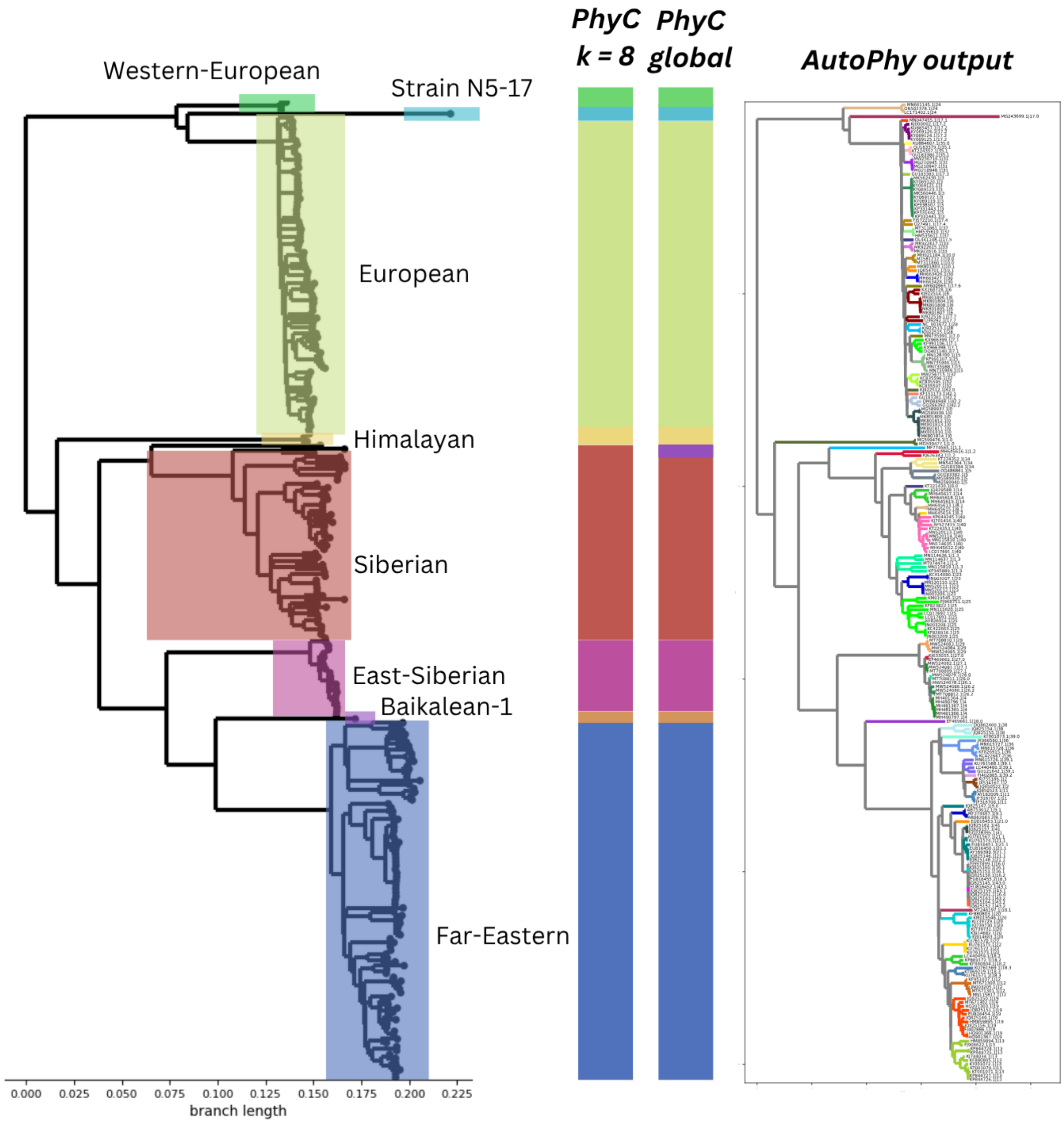


*Supplementary Figure 8: Tick-borne encephalitis viral phylogeny with annotated Nextstrain clades as well as PhytClust's chosen solution (k* = 9) and PhytClust's solution for k = 8 (left), and AutoPhy's output on the same tree (right). The AutoPhy visualization is reproduced directly from the software output.*

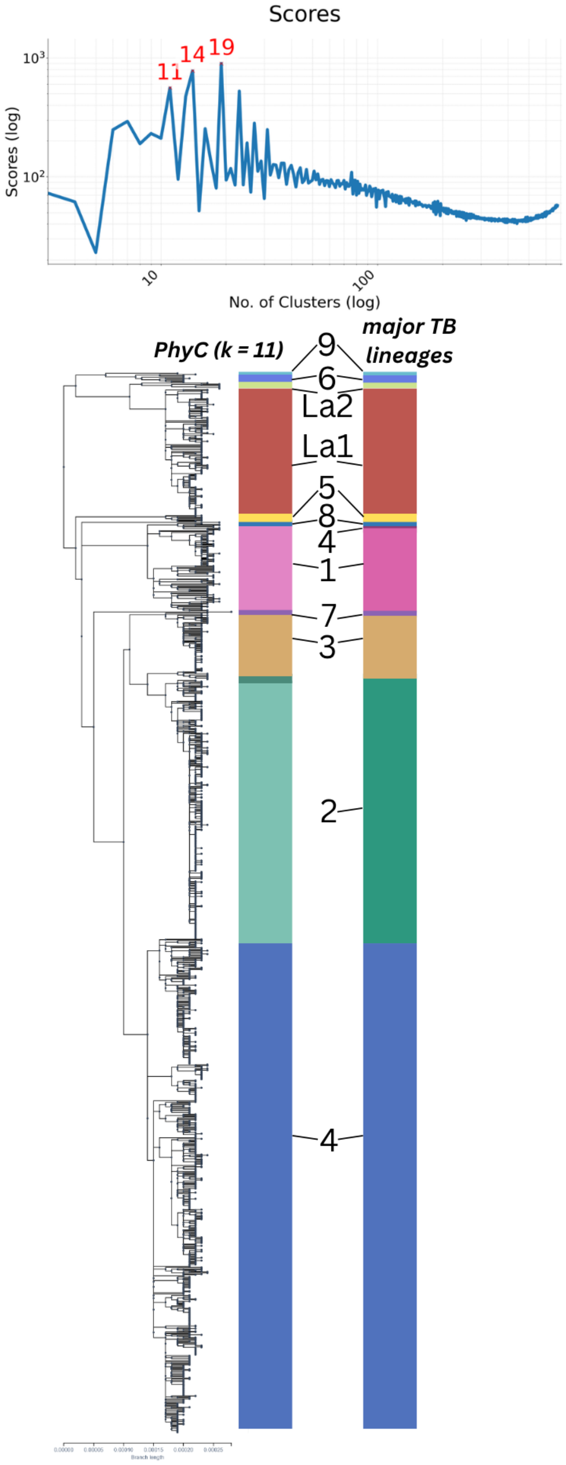


*Supplementary Figure 9: Tuberculosis phylogeny with the 11 major annotated lineages, alongside the PhytClust partition obtained for k = 11, and the corresponding PhytClust score profile. The PhytClust k = 11 solution agrees closely with the annotated lineage structure (V-measure = 0.98). Consistent with this, k = 11 appears as one of the five strongest candidate resolutions in the PhytClust validity index scores.*

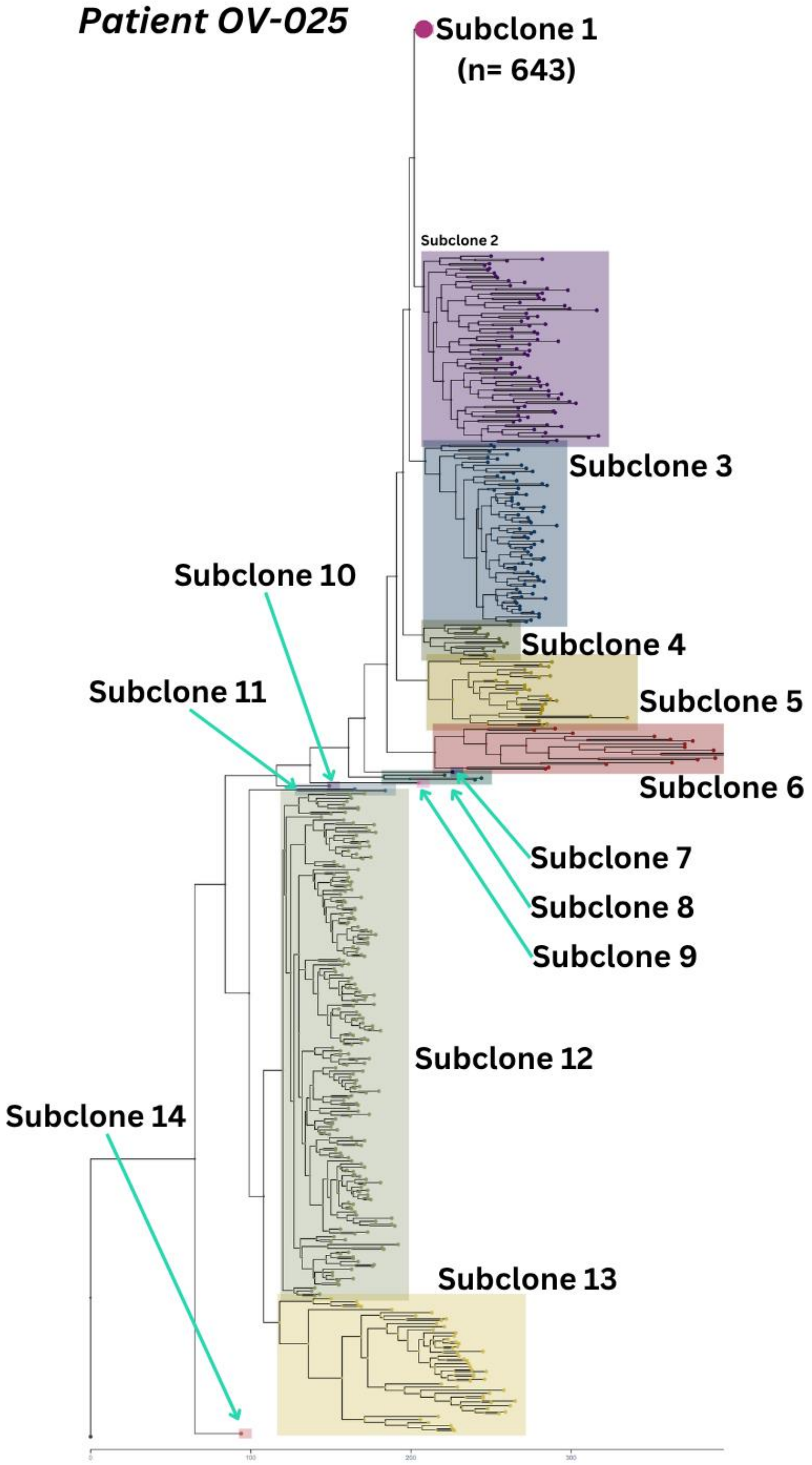


*Supplementary Figure 10: a) Complete OV-025 phylogeny with highlighted subclones identified by PhytClust. Subclone 1 with 643 cells was collapsed for ease of visualization.*

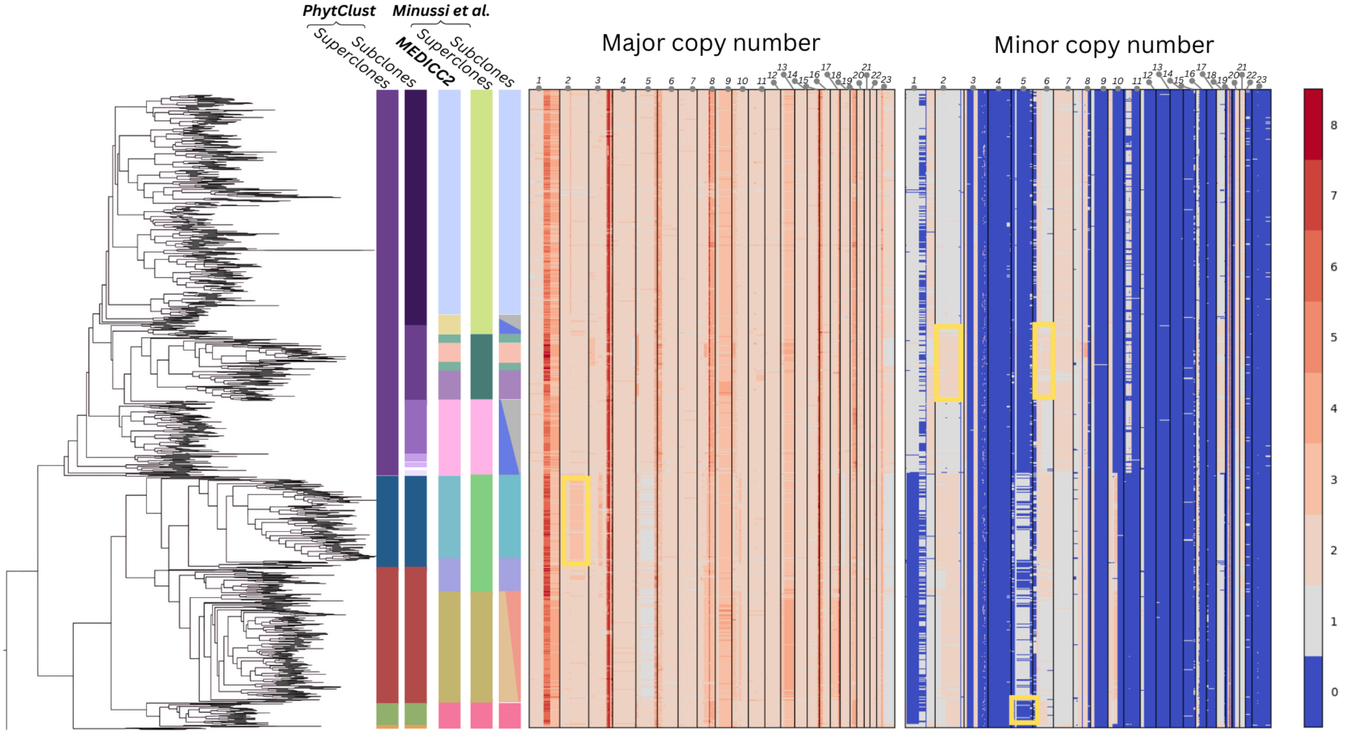


*Supplementary Figure 11: a) Minussi TN-2 phylogeny with original subclones and superclones from Minussi et al, subclones annotated by MEDICC2 authors as well as superclones (k = 6, $\alpha$ = 2.8) and subclones (k = 10, $\alpha$ = 2.5) identified by PhytClust.*

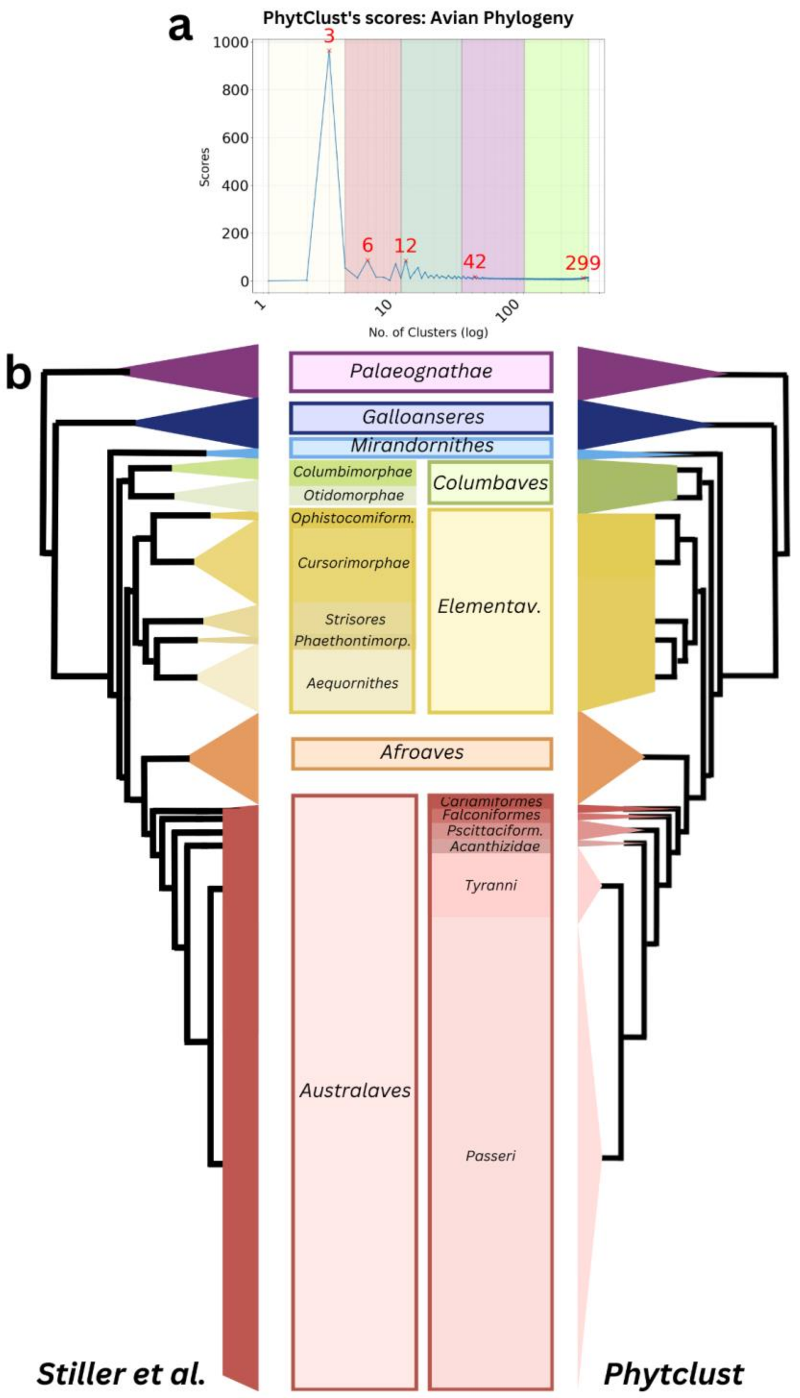


*Supplementary Figure 12: (a) PhytClust's scores for the avian phylogeny obtained from Stiller et al and stratified into 5 Clade Levels. (b) Comparison of clade annotations from Stiller et al and PhytClust's CL3 clusters.*

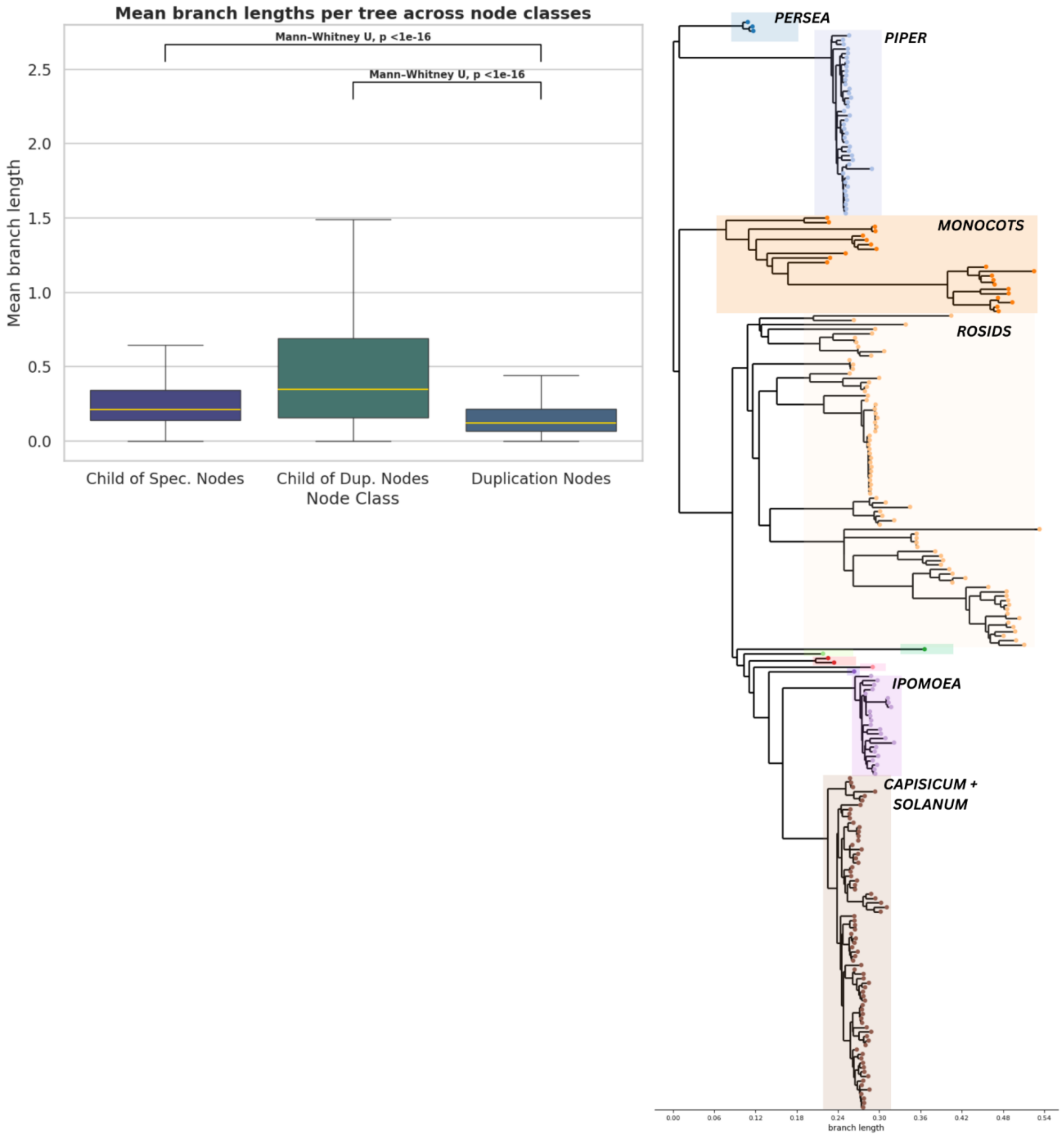


*Supplementary Figure 13: (a) distribution of z-normalised branch lengths of the phylogenetic trees from PANTHER database for all branches leading to a duplication node and branches following a duplication or speciation node.(b) Complete phylogenetic tree showing PhytClust cluster assignments, together with the most appropriate taxonomic annotations for each cluster*

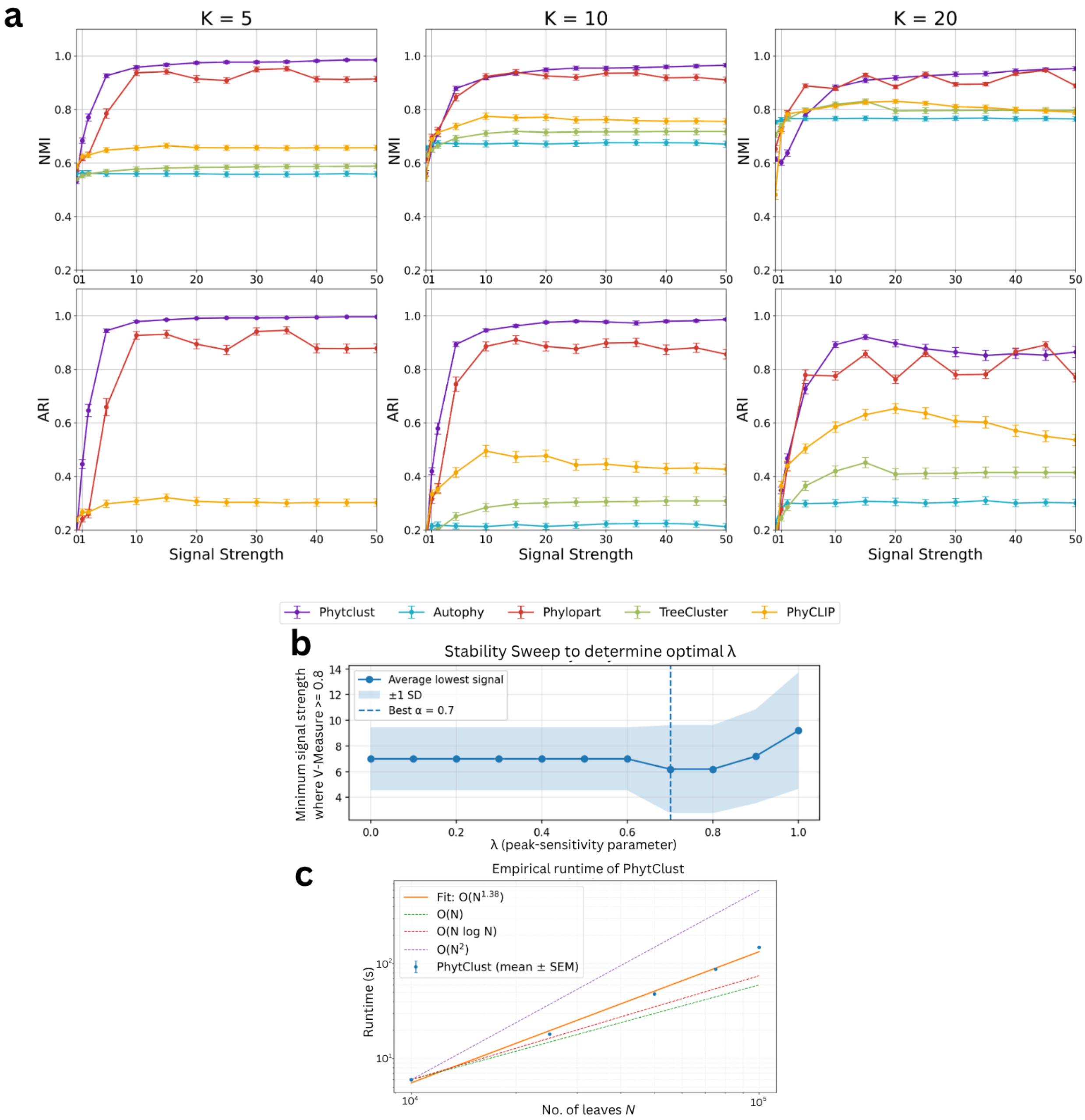


*Supplementary Figure 14: (a) Simulation results reporting Normalized Mutual Information (NMI) and Adjusted Rand Index (ARI) for PhytClust, AutoPhy, PhyCLIP, TreeCluster and PhyloPart of trees with N =100, k = 10. (b) Stability sweeps over various values of Lambda using simulated trees. (c) Empirical runtime scaling of PhytClust.*

### Pseudocode for Dynamic Programming:

```
Algorithm COMPUTE-DP-TABLE(pc)
V ← postorder(T)
for v in V do
   if v is leaf then
      dp_table[v][0] ← 0
      continue
   if v is a polytomy then
      (dp_table[v], polytomy_backptr[v]) ← COMPUTE-POLYTOMY-DP(v, pc)
      continue
   let ℓ, r be the children of v
   dp_table[v][0] ← dp_table[ℓ][0] + num_leaves[ℓ]·branch_length(ℓ)
              + dp_table[r][0] + num_leaves[r]·branch_length(r)

   if SUBTREE-ALL-ZERO(v) then
      continue

   for k ← 1 to num_leaves[v] do
      dp_table[v][k] ← min over i = 0 ... k-1 of
                  dp_table[ℓ][i] + dp_table[r][k-1-i]

      let i* be the minimiser
      backptr[v][0,k] ← i*
      backptr[v][1,k] ← k-1-i*

return (dp_table, backptr, polytomy_backptr)
```

```
Algorithm BACKTRACK(v, k, dp_table, backptr, polytomy_backptr)
    if v is leaf then
        return {v}
    if k = 0 then
        return {L(v)}                    ▷ all leaves of v form one cluster
    if v is a polytomy then
        return BACKTRACK-POLYTOMY(v, k, polytomy_backptr)
    let ℓ, r be the children of v
    i* ← backptr[v][0, k]
    j* ← backptr[v][1, k]
    return BACKTRACK(ℓ, i*, ...) ∪ BACKTRACK(r, j*, ...)
```

#### Proof A: Relationship Between the Dynamic Programming (β) Score and the Calinski–Harabasz Index

Aim: Given a phylogenetic tree with $N$ terminal nodes, find a score that when minimized, gives the most suitable clustering sets for $k$ monophyletic clusters. The DP score for a phylogenetic tree is given by:

$$\beta(c_k) = \sum_{c_i \in C_k} \sum_{i \in c_i} d(i, MRCA(i)) \quad (1)$$

Where,

$C_i$ is a given cluster in the set of clusters $C_k$

$C_k$ is any set of clusters possible for $k$ monophyletic clusters

*MRCA(i)* is the MRCA of the set of terminal nodes in a cluster

The Calinski Harabasz score is given by

$$\text{CH}(c_k) = \frac{\text{BSS}}{\text{WSS}} \text{X} \frac{\text{N-K}}{\text{K-1}}$$

*(2)*

Where,

*TSS* is the Total Sum of Squares

*BSS* is the between cluster Sum of Squares

*WSS* is the within cluster Sum of Squares

*N* is the number of data points

*k* is the number of clusters, for Euclidean distances

Since TSS = WSS + BSS,

$$CH(c_i\ ) = \frac{TSS - WSS}{WSS} \times\ m$$

where,

$$m = \frac{N-k}{k-1}$$

$$TSS = \sum_{i=1}^{N}(x_i - X)^2$$

$$WSS = \sum_{k=1}^{K}\sum_{c_i\, \epsilon\, C_k} ||x_i - C_k||^2$$

$x_i$ are data points in a cluster, X is the centroid of a cluster.

Patristic distances in a phylogenetic tree are equivalent to squared Euclidean distances, as proven by de Vienne et al[67]. If we consider the MRCA of terminal nodes in a cluster to be the local centroid and the root to be the global centroid:

$$\beta(1) = \sum_{c_i\, \epsilon\, C_k}\sum_{i\, \epsilon\, c_i} d(i, \rho) = TSS$$

Where ρ is the root of the phylogenetic tree.

Similarly,

$$\beta(c_k) = \sum_{c_i \, \epsilon \, C_k} \sum_{i \, \epsilon \, c_i} d(i, MRCA(i)) \ = WSS$$

The Calinski Harbasz score can now be rewritten as:

$$\mathrm{CH}(c_k) = \frac{\beta(1) - \ \beta(c_k)}{\beta(c_k)} \mathrm{X} \ m$$

∴

$$1/\mathrm{CH}(c_k) = \frac{\beta(c_k)}{m(\beta(1) - \ \beta(c_k))}$$

As $\mathrm{CH}(c_k)$ increases, $\beta(c_k)$ decreases. If we consider $\mathrm{CH}(c_k) = y$,,$\beta(1)$ = a, $\beta(c_k)$ = x, we can rewrite equation as:

$$\frac{1}{x} = \frac{y}{m(x-a)}$$

$$1 \ = \ y\left(\frac{x}{m}\right) - y\left(\frac{x^2}{m}\right)$$

Hence, the relationship between both scores is described by a quadratic function. The graph for this relationship would be parabolic in nature.
